# Sequential regulation of maternal mRNAs through a conserved cis-acting element in their 3’UTRs

**DOI:** 10.1101/285569

**Authors:** Pooja Flora, Siu Wah Wong-Deyrup, Elliot Todd Martin, Ryan J Palumbo, Mohamad Nasrallah, Andrew Oligney, Patrick Blatt, Dhruv Patel, Gabriele Fuchs, Prashanth Rangan

## Abstract

Maternal mRNAs are synthesized during oogenesis to initiate the development of future generations. Some maternal mRNAs are determinants of somatic or germline fate and must be translationally repressed until embryogenesis. However, the translational repressors themselves are also temporally regulated. We use *polar granule component* (*pgc*), a *Drosophila* maternal mRNA, as a model system to ask how maternal mRNAs are repressed while the regulatory landscape is continually shifting. *pgc*, a potent transcriptional silencer and germline determinant, is translationally regulated throughout oogenesis. We find that the 3’UTR of *pgc* mRNA contains a conserved ten-nucleotide sequence that is bound by different conserved RNA binding proteins (RBPs) at different stages of oogenesis to continuously repress translation except for a brief expression in the stem cell daughter. Pumilio (Pum) binds to this sequence in undifferentiated and early differentiating oocytes and recruits other temporally restricted translational regulators to block *pgc* translation. After differentiation, Pum levels diminish and Bruno (Bru) levels increase, allowing Bru to bind the same 3’UTR sequence and take over translational repression of *pgc* mRNA. We have identified a class of maternal mRNAs regulated during oogenesis by both Pum and Bru, including *Zelda*, activator of the zygotic genome, which contain this core 10-nt regulatory sequence. Our data suggests that this hand off mechanism is more generally utilized to inhibit translation of maternal mRNAs during oogenesis.

## Introduction

The germ line gives rise to the eggs and sperm that launch the next generation. Upon fertilization, the egg differentiates into every cell lineage present in the adult organism, including a new germ line and is therefore totipotent (Seydoux and Braun 2006; Cinalli et al. 2008). Pivotal to the egg’s task of kick-starting the next generation is a maternally synthesized trust fund of mRNAs that are deposited into the egg during oogenesis (Lasko 2012). Post fertilization, and prior to zygotic genome activation, translation of these maternally supplied mRNAs help power early development (Evans 2005; Zhang and Smith 2015; Tadros and Lipshitz 2009; Becalska and Gavis 2009; Lee et al. 2014). Some of the maternally supplied mRNAs code for key determinants of both somatic and germ cell fate, and thus need to be exquisitely regulated both during oogenesis and early embryogenesis.

RNA binding proteins (RBPs) regulate the maternal pool of mRNA through interactions with specific sequences within the 3’ untranslated regions (UTRs) of their target mRNAs (Kuersten and Goodwin 2003; Rosario et al. 2017; Moor et al. 2005; Slaidina and Lehmann 2014; Johnstone and Lasko 2001; Evans 2005). Loss of RBPs during oogenesis results in death, sterility or germ line to soma trans-differentiation (Ciosk 2006; Forbes and Lehmann 1998; Mak et al. 2016; Tsuda 2003). This suggests that RBPs are critical for silencing key somatic and germ line determinants during oogenesis. Consistent with this observation, it has been shown that gene regulation during oogenesis and early embryogenesis relies primarily on the 3’UTRs of mRNAs rather than on their (Merritt et al. 2008; Rangan et al. 2009a). Additionally, loss of specific sequences in the 3’UTR of maternal mRNAs results in their dysregulation (Kim-Ha et al. 1995; Wharton and Struhl 1991). However, several RBPs that are regulators of translation also fluctuate in levels, with these fluctuations promoting critical developmental transitions. For example, during *C. elegans* oogenesis two RBPs, GLD-1 and MEX-3, whose loss results in germ line to soma trans-differentiation, have a reciprocal expression pattern (Jones et al. 1996; Draper et al. 1996; Mootz et al. 2004; Ciosk 2006). In human fetal ovary, RBPs such as Deleted in Azoospermia-like (DAZL) play an important role in regulating RNA targets, such as *TEX11*, a gene required for recombination and DNA repair, via its 3’UTR (Rosario et al. 2017). However, DAZL itself has a dynamic expression pattern during human oogenesis in which it is robustly expressed in the pre-meiotic and post-meiotic germ cells but absent during meiotic stages (Anderson et al. 2007; He et al. 2013). The conundrum remains as to how mRNAs can be continually silenced during oogenesis when the RBPs that regulate them fluctuate.

*Drosophila* oogenesis is an excellent model to investigate how maternal mRNAs are continuously regulated. Oogenesis in *Drosophila* begins when germline stem cells (GSCs) divide to both self-renew and give rise to a stem cell daughter called a cystoblast (CB) (Fig. 1A-B) (Chen and McKearin 2003). The CB differentiates by undergoing four incomplete mitotic divisions to give rise to a 2-, 4-, 8-, and 16-cell cyst (Fig. 1B) (Koch et al. 1967; McKearin and Ohlstein 1995; McKearin and Spradling 1990). Of these sixteen cells, one is designated as the oocyte and the other fifteen cells become nurse cells (Fig. 1A) (Spradling et al. 1997); the maternal mRNAs and proteins synthesized by the nurse cells are deposited into the oocyte (Zalokar 1960). The oocyte and surrounding nurse cells are encapsulated by somatic cells to form an egg chamber, which progresses through successive developmental stages (Margolis and Spradling 1995; Gilboa and Lehmann 2004a). These maternal mRNAs which are deposited into the oocyte need to be post-transcriptionally regulated to promote proper oogenesis and embryogenesis (Evans 2005; Richter and Lasko 2011; Lasko 2012; Laver et al. 2015).

**Figure 1.**
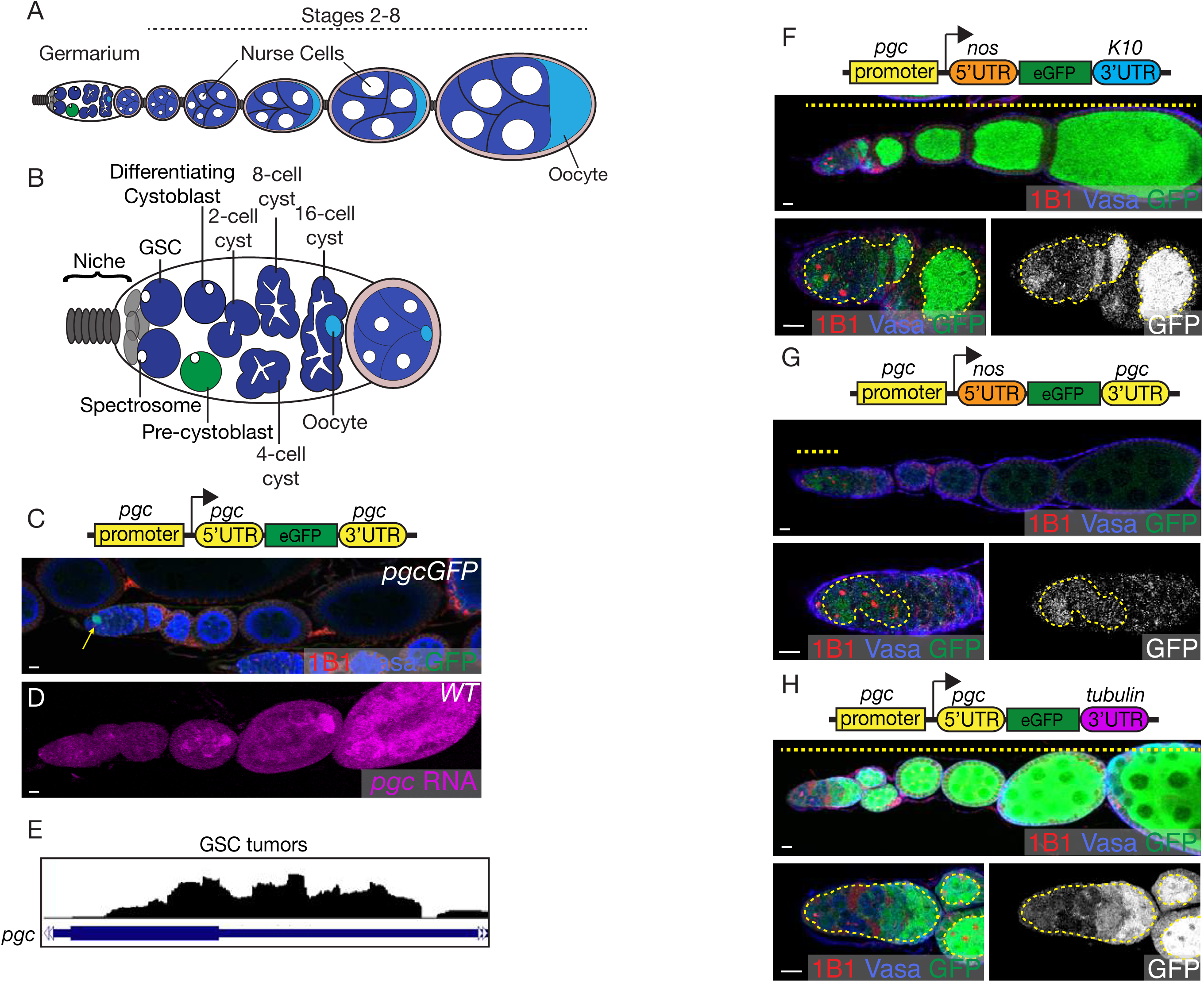
Pgc is translationally regulated via its UTRs. (A) Schematic representation of a female *Drosophila* ovariole. The *Drosophila* ovary is composed of 16-18 ovarioles, each of which is an assembly line of egg chambers at 14 different stages. Each chamber is encapsulated by somatic follicle cells and contains fifteen nurse cells that provide mRNAs and proteins to the developing oocyte. (B) A schematic representation of a germarium that is present at the anterior most tip of the ovariole. The germline stem cells (GSCs) marked in blue are supported and maintained by the somatic niche marked in gray. Each GSC divides asymmetrically to give rise to a pre-cystoblast (pre-CB), shown in green. The pre-CB then turns on a differentiation factor, *bag-of-marbles* (*bam*) and undergoes four incomplete mitotic divisions to give rise to a 16-cell cyst. The single cells of the germarium can be identified by the circular structure called the spectrosome and the differentiating cysts can be identified by the branched structures called fusomes. One of the cells from the 16-cell cyst becomes the oocyte (light blue) while the other fifteen become the nurse cells. (C) The ovariole of a transgenic fly created by fusing GFP to the *pgc* 5’ and 3’UTR under the control of the *pgc* promoter was stained with 1B1 (red), Vasa (blue) and GFP (green). Expression of GFP is restricted to the pre-CB in the germarium. (D) The ovariole of a wild-type fly probed for *pgc* RNA (magenta) using FISH, shows that *pgc* RNA is present throughout oogenesis, with increasing levels being deposited in the developing oocyte. (E) RNA-seq track of *pgc* in *nos-GAL4*>*UAS-tkv* ovaries show *pgc* RNA is transcribed in the GSCs. (F) The ovariole of a transgenic fly created by fusing GFP to the *nos* 5’ and *K10* 3’UTR under the control of the *pgc* promoter was stained with 1B1 (red), Vasa (blue) and GFP (green). GFP expression shows that the promoter is active in the GSCs. (G) The ovariole of a transgenic fly created by fusing GFP to *nos* 5’ and *pgc* 3’UTR under the control of the *pgc* promoter was stained with 1B1 (red), Vasa (blue) and GFP (green). There is a loss of GFP regulation only in the earliest stages of oogenesis. (H) The ovariole of a transgenic fly created by fusing GFP to the *pgc* 5’ and *tub* 3’UTR and under the control of the *pgc* promoter was stained with 1B1 (red), Vasa (blue) and GFP (green). There is a loss of GFP regulation throughout oogenesis, including at the earliest stages. Scale bars: 10μm.

*Polar granule component* (*pgc*) is a superb candidate to address how such maternal mRNAs are regulated during the developmental transitions of oogenesis. *pgc* is synthesized during oogenesis and provided to the oocyte, where it localizes to the germ plasm (Nakamura et al. 1996). While *pgc* mRNA is continuously present, Pgc is only translated in two short pulses; once in the CB during oogenesis and once in the germ cells during embryogenesis (Flora et al. 2018; Hanyu-Nakamura et al. 2008). Pgc expression in the CB is required to promote the cell’s timely differentiation (Flora et al. 2018). Pgc expression in the germ cells is required to repress the expression of somatic genes which could interfere with germ line specification (Hanyu-Nakamura et al. 2008). Pgc performs these tasks by causing global transcriptional silencing through targeting the basal transcriptional elongation machinery of RNA polymerase II (Hanyu-Nakamura et al. 2008; Flora et al. 2018; Martinho et al. 2004). *pgc* can even suppress transcription in other cell types upon ectopic expression (Timinszky et al. 2008). The strong effects of Pgc on transcription lead to a requirement for strict regulation of *pgc* translation in the cells in which it is normally found. It is known that the 3’UTR of *pgc* mRNA is sufficient to mediate such translational control after differentiation (Rangan et al. 2009b); however it is currently not known if *pgc* is regulated transcriptionally or translationally prior to differentiation as well as what trans-acting factors bind to its 3’UTR after differentiation.

Temporally restricted RBPs that bind to 3’UTRs regulate critical developmental transitions during *Drosophila* oogenesis by controlling translation of their targets. Pumilio (Pum), an RBP that belongs to the conserved <u>P</u>um and <u>F</u>em-3 binding factor (PUF) domain family of proteins, is present at high levels in the undifferentiated cells in the ovary including GSCs, CBs and early differentiating cysts (Zhang et al. 1997; Carreira-Rosario et al. 2016; Lin and Spradling 1997; Forbes and Lehmann 1998; Wickens et al. 2002). Pum represses translation of differentiation-promoting mRNAs in the GSCs thereby preventing stem cell loss (Forbes and Lehmann 1998; Joly et al. 2013). Pum expression is attenuated in the differentiated stages allowing for the expression of the differentiation promoting mRNAs (Forbes and Lehmann 1998; Carreira-Rosario et al. 2016). *Drosophila* Bruno 1 (Bru), a <u>C</u>UGBP and <u>E</u>TR-3 <u>L</u>ike <u>F</u>actor (CELF) superfamily protein, is expressed at increasing levels during differentiation and is then maintained for the rest of oogenesis (Xin et al. 2013; Sugimura and Lilly 2006; Webster et al. 1997). Bru regulates several maternal mRNAs post-differentiation during oogenesis (Good et al. 2000; Moraes et al. 2006; Schüpbach and Wieschaus 1991; Webster et al. 1997; Snee et al. 2014; Filardo and Ephrussi 2003; Moore et al. 2009; Castagnetti et al. 2000). Thus, Pum and Bru have reciprocal temporal regimes and thus could act jointly to repress targets throughout oogenesis. However, it is not known whether further repression is required of Pum targets after differentiation, or Bru targets prior to differentiation.

Pum and Bru during their regimes can use various cofactors to mediate translational repression using distinct mechanisms. Pum partners with Nanos (Nos) in the GSCs to recruit translation modulators such as Twin, a deadenylase causing a shortening of the poly(A)-tail (Joly et al. 2013). Pum can also recruit Brain Tumor (Brat) which is known to modulate translation by interacting with *<u>D</u>rosophila* Eukaryotic translation initiation factor <u>4E</u> <u>H</u>omologous <u>P</u>rotein (d4EHP), a cap binding protein (Cho et al. 2006; Harris et al. 2011). Bru can form oligomers to form silencing particles or can partner with Cup, which associates with the 5′-cap binding initiation factor eIF4E, to regulate mRNAs (Nakamura et al. 2004; Kim-Ha et al. 1995; Chekulaeva et al. 2006; Filardo and Ephrussi 2003; Kim et al. 2015b). Why certain mechanisms are preferred over others at a particular temporal regime is not known.

Here, we elucidate an intricate and elegant control mechanism ensuring handoff of translational repression of a germ line determinant, *pgc*, from one set of regulators to another, with the exception of a single gap in the CB. This governs the critical temporal control of *pgc* production just in CBs, ensuring proper maintenance of GSCs and their conversion into differentiated progeny. We demonstrate that this control depends on a 10-nucleotide (nt) sequence in the 3’UTR of *pgc* mRNA. In the undifferentiated stages, we find that Pum binds the 10-nt sequence and partners with Nos and the CCR4-Not complex to regulate *pgc* mRNA in a poly(A) dependent manner. When Nos levels drop in CBs, *pgc* is expressed. After CB differentiation, Pum switches partners to use Brat to suppress *pgc* in the early differentiating cysts in a cap dependent manner. However, when Pum levels diminish, *pgc* mRNA is bound by Bru via the same 10-nt sequence to translationally regulate it. Bru recruits Cup to silence *pgc* translation also in a cap dependent manner. We find that a class of maternal mRNAs, including *zelda*, which play pivotal roles during development, are also regulated by both Pum and Bru and contain this 10-nucleotide sequence. This suggests that the sequential hand off of mRNAs between Pum and Bru is broadly utilized to control fine-scale translation of maternal RNAs. We propose that this handoff mechanism from one set of trans-acting factors that utilizes a poly(A) shortening to another set of trans-acting factors that utilizes cap dependent mechanism is required to protect mRNAs post-differentiation and prime them for translation during embryogenesis.

## Results

### Pgc is translationally regulated via its UTRs

During oogenesis, Pgc protein is expressed only in the CBs, where it promotes timely differentiation (Fig. 1C) (Flora et al. 2018). To assess if this temporal specificity of Pgc protein production is due to transcriptional or translational regulation, we first carried out fluorescent *in situ* hybridization (FISH) in both wild-type and *pgcGFP* reporter ovaries. *pgc* transcription in the GSCs was difficult to discern because of the low resolution of FISH in the germarium, however we did detect *pgc* mRNA in all later differentiated stages (Fig. 2D, Supplemental Fig. S1A-C). To assess *pgc* mRNA expression in the GSCs through an alternate method, we over-expressed the self-renewal signaling receptor, Thick Veins Receptor (TKV), to enrich for GSCs and then sequenced their transcriptome (Xie and Spradling 1998). We detected 88 transcripts per million (TPM) of *pgc* transcript indicating that the mRNA is transcribed in the GSCs (Fig. 1E, Supplemental Fig. S1D). To further substantiate that the *pgc* promoter is active in the GSCs, we created a reporter construct in which the *pgc* promoter drives the expression of GFP flanked by the *nos* 5’UTR and *K10* 3’UTR, which are not translationally silenced during oogenesis (Fig. 1F) (Serano et al. 1994; Gavis and Lehmann 1994; 1992). We observed GFP expression throughout oogenesis, including in the GSCs. This suggests that the maternal *pgc* mRNA is transcribed from the GSCs onward throughout oogenesis and is under strict translational regulation pre- and post-differentiation (Rangan et al. 2009b).

**Figure 2.**
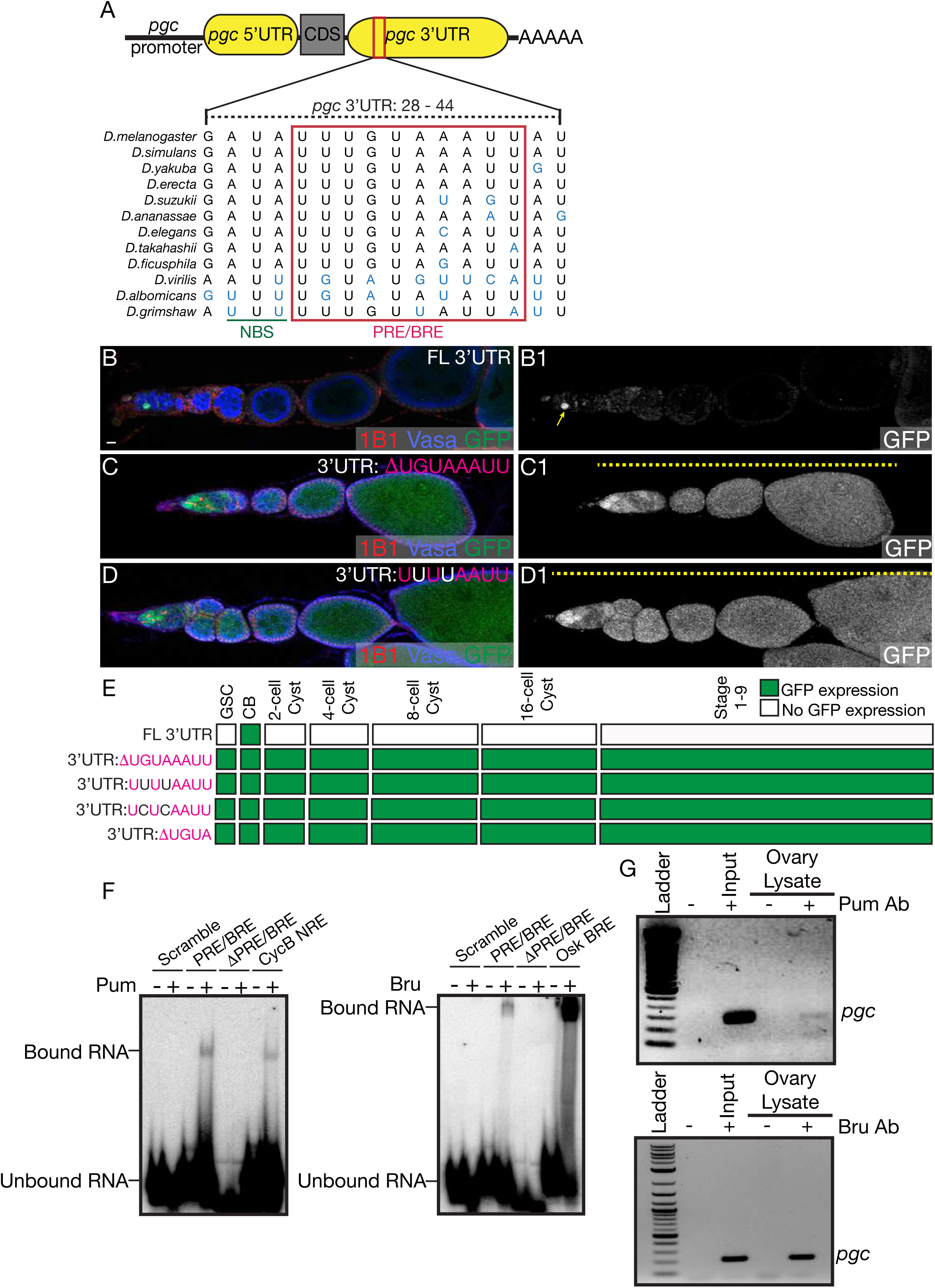
A cis-element in the *pgc* 3’UTR that binds both Pum and Bru is required for translational control throughout oogenesis. (A) The NBS and PRE/BRE sequence identified in the *pgc* 3’UTR of *Drosophila melanogaster* that can bind Pum and Bru is conserved in 12 species of Drosophilids. (B) The ovariole of a transgenic fly created by fusing GFP to the *pgc* 5’ and full length (FL) 3’UTR under the control of the *pgc* promoter was stained with 1B1 (red), Vasa (blue) and GFP (green). GFP reporter and thus normal Pgc expression was restricted to the pre-CB. (C) The ovariole of a transgenic fly created by fusing GFP to the *pgc* 5’ and PRE/BRE sequence deleted 3’UTR (3’UTR:ΔUGUAAAUU) under the control of the *pgc* promoter was stained with 1B1 (red), Vasa (blue) and GFP (green). A loss of GFP regulation was observed throughout oogenesis in the absence of the PRE/BRE sequence. (D) The ovariole of a transgenic fly created by fusing GFP to the *pgc* 5’ and 3’UTR (3’UTR: UUUUAAUU) where the UGUA core motif was mutated to UUUU and driven under the control of the *pgc* promoter was stained with 1B1 (red), Vasa (blue) and GFP (green). A loss of GFP regulation was observed throughout oogenesis when the UGUA sequence in the PRE was mutated to UUUU. (E) A developmental profile of GFP expression in different stages of oogenesis in transgenes where the PRE sequence was either deleted or mutated (3’UTR:ΔUGUAAAUU, 3’UTR: UUUUAAUU, 3’UTR: UCUCAAUU and 3’UTR: ΔUGUA) compared to FL 3’UTR. (F) EMSA shows that purified recombinant RNA binding domain of Pum protein binds to the PRE/BRE of *pgc* 3’UTR sequence *in vitro*. A scrambled RNA sequence shows no binding while the NRE sequence from the *CycB* 3’UTR shows binding. EMSA shows that purified full length recombinant Bru protein binds to the PRE/BRE sequence of *pgc* 3’UTR *in vitro*. A scrambled RNA sequence shows no binding while the BRE sequence from the *Osk* 3’UTR shows binding. (G) RT-PCR of *pgc* carried out on RNA samples extracted after an IP experiment with Pum antibody and Bru antibody in wild-type ovary lysate, respectively show that *pgc* RNA associates with Pum and Bru *in vivo*. Scale bars: 10μm.

Given that 5’UTR and 3’UTR of an mRNA are commonly recognized by sequence-specific RBPs that regulate translation (Wilkie et al. 2003), we wanted to test the potential role of both the 5’ and 3’UTR of *pgc* in repressing translation in the GSCs. *pgc* mRNA has two annotated 5’UTRs; to determine which one was specifically expressed in the GSCs, we designed primers that distinguish these two forms. We carried out PCR on RNA enriched from GSCs by over-expressing the self-renewal signaling receptor, TKV, and for CBs, by using a mutation for differentiation factor, *bam* (Xie and Spradling 1998; McKearin and Ohlstein 1995). We found that only the short form was expressed in the GSCs and CBs (Supplemental Fig. S1E). To determine if this short *pgc* 5’UTR is required for translational regulation of *pgc*, we swapped it with the *nos* 5’UTR in a GFP reporter construct that still retained the *pgc* 3’UTR and the *pgc* promoter. We found that the absence of the *pgc* 5’UTR results in an upregulation of GFP protein expression in the GSCs but not in later stages (Fig. 1G). Our results indicate that in GSCs, the *pgc* 5’UTR is required for translational regulation, while the 3’UTR is not sufficient (Fig.1G). In differentiated stages, the 3’UTR alone is sufficient to mediate translational regulation (Fig.1G). To test if the 5’UTR is sufficient for translational regulation in GSCs, we created a construct with the *pgc* 5’UTR and non-repressed *tubulin* (*tub*) 3’UTR flanking GFP under the control of the *pgc* promoter (Fig. 1H). GFP was still expressed in the GSCs as well as in later differentiating stages and egg chambers demonstrating that the 5’UTR alone is not sufficient for translational regulation (Fig. 1H). Taken together, we conclude that both the *pgc* 5’ and 3’UTR are required for translational control pre-differentiation in the GSCs, and that the 3’UTR alone is sufficient post-differentiation in the cysts and egg chambers.

### A cis-element in the *pgc* 3’UTR that binds both Pum and Bru is required for translational control throughout oogenesis

We predicted that cis-acting sequences in either the 5’ or 3’ UTRs of *pgc* could regulate translation during oogenesis by recruiting trans-acting factors such as RBPs. To identify these sequences, we carried out a phylogenetic analysis of the *pgc* 5’ and 3’UTR in Drosophilids that were separated by 40 million years of evolution and discovered several regions of conservation in the 3’UTR (Supplemental Fig. S2A). We could not identify unique conserved regions in the *pgc* 5’UTR as the sequence overlaps with the coding region of *Type III alcohol dehydrogenase* (*T3dh*) on the opposite chromosomal arm. We also used algorithms that search for RBP binding sequences, and did not find any in the short form 5’UTR of *pgc* (Bailey et al. 2009). In the 3’UTR, a conserved 10-nt sequence, UUUGUAAAUU, stood out (Fig. 2A, Supplemental Fig. S2A). This sequence closely matches the sequences AUUGUACAUA and UUUGUAAUUU, which have been previously described as a the <u>P</u>umilio <u>R</u>esponse <u>E</u>lement (PRE), which is part of the <u>N</u>anos <u>R</u>esponse <u>E</u>lement (NRE) in *hunchback* (*hb*) and *Cyclin B* (*CycB*), respectively (Wharton and Struhl 1991; Weidmann et al. 2016; Murata and Wharton 1995; Kadyrova et al. 2007). PREs are known to bind Pum, which then recruits Nos, to the bind to the <u>N</u>anos <u>B</u>inding <u>S</u>equence (NBS) resulting in translational regulation of RNAs (Fig. 2A) (Asaoka-Taguchi et al. 1999; Kadyrova et al. 2007; Muraro et al. 2008; Sonoda and Wharton 1999; Murata and Wharton 1995). This sequence in the *pgc* 3’UTR can also bind another conserved RBP, Bru. Pum binds to the UGUA motif while Bru binds to a uU^G/A^U^G/A^U^G/A^Uu motif which is described as the <u>B</u>runo <u>R</u>esponse <u>E</u>lement (BRE) (Kim-Ha et al. 1995; Wharton and Struhl 1991).

We asked if this conserved 10-nt sequence that is predicted to bind two RBPs can regulate *pgc* translation. To test this, we generated a reporter construct that deleted 8-nt of the conserved sequence including the UGUA motif that is known to bind Pum and the uU^G/A^U^G/A^ motif that binds Bru. This resulted in an upregulation of translation throughout oogenesis (Fig. 2B-C, E Supplemental Fig. S2D). We also generated three other transgenes in which we mutated the core UGUA motif to UUUU and UCUC and also deleted the core UGUA motif respectively. We found that all three of these changes resulted in ectopic GFP expression throughout oogenesis (Fig. 2D-E, Supplemental Fig. S2B-D). Thus, we conclude that the conserved 10-nt sequence in the *pgc* 3’UTR that is predicted to bind Pum and Bru controls translation of *pgc* throughout oogenesis.

To determine if the conserved sequence actually binds Pum and Bru as predicted, we purified the recombinant RNA binding domain of Pum (residues 1091-1426) and full length Bru and carried out Electrophoresis Mobility Shift Assay (EMSA) experiments (Supplemental Fig. S2E) (Chekulaeva et al. 2006; Weidmann et al. 2016). As positive controls, we utilized the NRE in *CycB* and the BRE in *Oskar’s* (*osk*) 3’UTR and first demonstrated that our recombinant Pum and Bru bound the NRE and BRE, respectively (Fig. 2F) (Kadyrova et al. 2007; Kim-Ha et al. 1995). Both Pum and Bru also bound the PRE in the 3’UTR of *pgc*, but only in the presence of the conserved 10-nt sequence (Fig. 2F). To test, if Pum and Bru also bind to *pgc* mRNA *in vivo*, we performed an immuno-precipitation (IP) experiment with anti-Pum antibody and with anti-Bru antibody in wild-type ovary lysates. We observed that *pgc* mRNA associated with both Pum and Bru upon their respective pull down (Fig. 2G, Supplemental Fig. S2F). Thus, we conclude that Pum and Bru bind to the 10-nt PRE of *pgc* 3’UTR *in vitro* and to *pgc* mRNA *in vivo*.

### Pum and its co-factor Nos regulate Pgc translation in the GSCs and early differentiating cysts

We asked if *pgc* was translationally regulated by Pum and Bru during oogenesis, and in particular, given their inverse expression patterns, if they might each govern distinct phases. Pum is expressed from the GSCs to the 8-cell cysts and is attenuated from the 16-cell cyst onwards (Supplemental Fig. S2G-G2’) (Forbes and Lehmann 1998; Carreira-Rosario et al. 2016). Bru levels are low from GCSs to the 8-cell cyst stage, but are high in the 16-cell cyst stage and throughout later oogenesis (Supplemental Fig. S2G-G2’) (Xin et al. 2013; Sugimura and Lilly 2006; Webster et al. 1997). Thus, we hypothesized that Pum may regulate *pgc* translation until the 8-cell cyst and Bru thereafter. We first focused on Pum and its potential role in regulating *pgc* translation during early oogenesis. Pum requires co-factors to regulate translation and can use distinct partners and thus multiple mechanisms. Pum is known to recruit Nos and Twin, a deadenylase, to NRE-containing 3’ UTRs to induce poly(A)-tail shortening in *Drosophila* embryonic germ cells (Sonoda and Wharton 1999; Kadyrova et al. 2007). During oogenesis Twin is ubiquitously expressed (Temme et al. 2010; Joly et al. 2013) and Nos protein is present in all stages, except for in the pre-CB where Pgc is expressed (Supplemental Fig. S3A-B1) (Forbes and Lehmann 1998; Li et al. 2009). We therefore hypothesized that Pum, might be regulating Pgc expression with Nos and Twin only until the cyst stages, during which time a drop in Nos expression in the pre-CBs would allow for Pgc expression there.

To test this hypothesis, we separately assayed for PgcGFP expression in *pum*, *nos* and *twin* mutants. We observed that in the absence of each of these genes, the reporter was ectopically expressed in the GSCs, as marked by pMAD, and in 2- and 4-cell cysts (Fig. 3A-D1, Supplemental Fig. S3C-F). Ectopic expression in the GSCs was also observed upon germline depletion of *pum, nos* and *twin* via RNAi (Supplemental Fig. S3G-I, L). Twin is a deadenylase and is part of the CCR4-Not complex (Morris 2005; Temme et al. 2010; Chicoine et al. 2007; Temme et al. 2014; Fu et al. 2015). To determine if other members of this complex were also involved in regulating the *pgc* 3’UTR, we depleted Pop2 and Not1 in the germ line using RNAi and assayed for GFP expression. Compared to *pgcGFP*, depletion of Pop2 and Not1 resulted in ectopic expression of the reporter from the GSCs to the 4-cell cysts consistent with what we observed in the *nos*, *pum*, and *twin* mutants (Supplemental Fig. S3J-L). We also observed that loss of *pum* and *twin* results in an elevated GFP expression in the 8-cell cyst. Based on these immunofluorescence (IF) experiments, we generated a developmental profile to show the temporal loss of translational regulation of GFP at each stage of development in *pum, nos and twin* when compared to control *pgcGFP* ovarioles (Fig. 3E). Taken together we can conclude that *pgc* is regulated by Nos, Pum and Twin from GSCs to the 4-cell cyst stage via the CCR4-Not complex. In the pre-CB, when Nos is absent, Pgc is expressed even though Pum and Twin proteins are still present. This suggests that Pum and Twin alone are not sufficient for regulating *pgc* in the pre-CB and require the presence of their co-regulator Nos.

**Figure 3.**
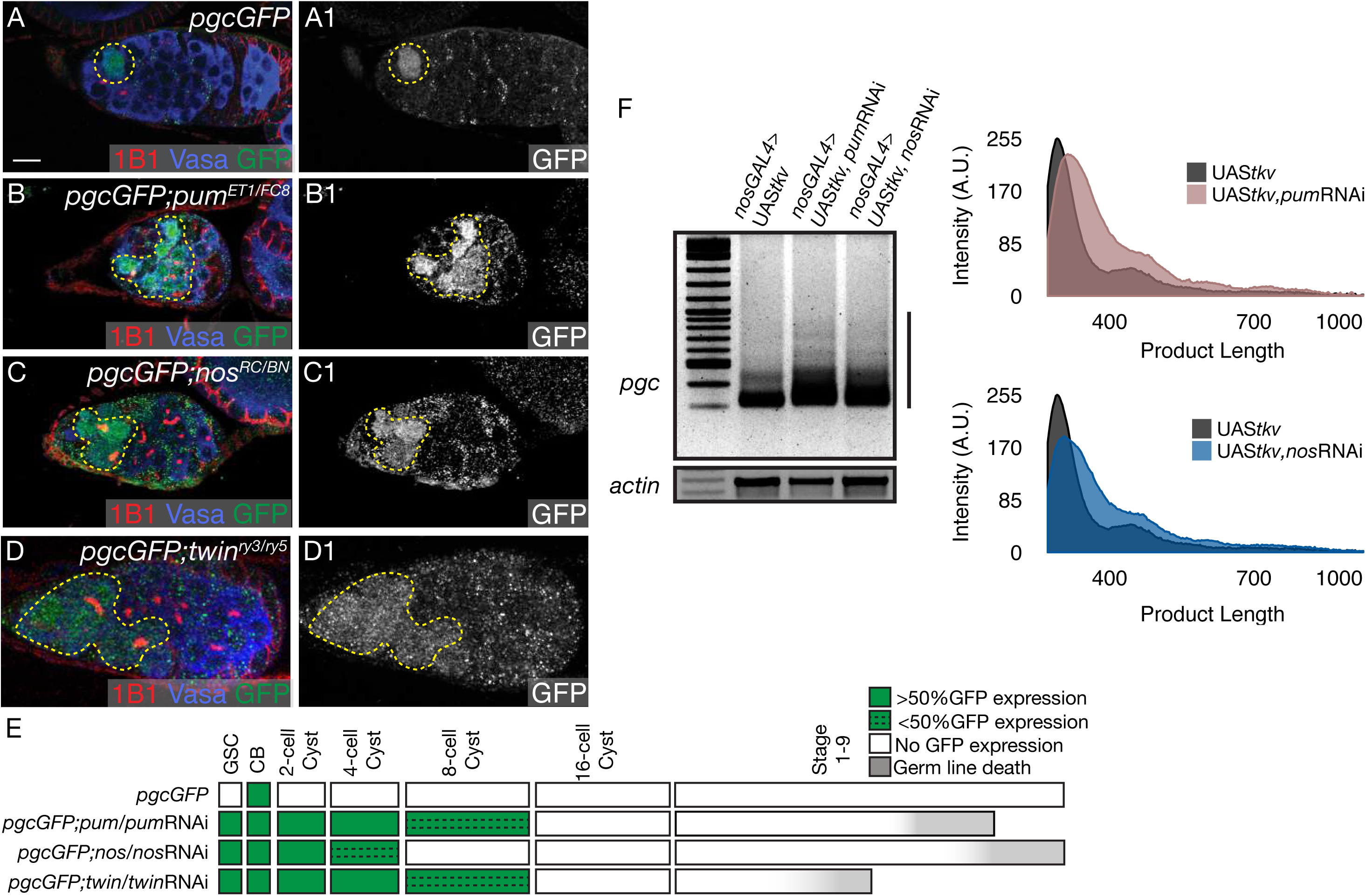
Pum and its co-factor Nos regulate Pgc translation in the GSCs and early differentiating cysts. (A, A1) The germarium of a *pgcGFP* ovary stained with 1B1 (red), Vasa (blue) and GFP (green) shows expression of GFP only in the pre-CB. GFP channel shown in gray scale in A1. (B, B1) The germarium of a *pgcGFP; pum*^*ET1/FC8*^ mutant ovary stained with 1B1 (red), Vasa (blue) and GFP (green) shows aberrant expression of GFP from the GSCs to the 8-cell cyst (100% from GSC to 4-cell cyst, 32% in 8-cell cyst, n= 25 germaria). GFP channel shown in gray scale in B1. (C, C1) The germarium of a *pgcGFP; nos*^*RC/BN*^ mutant ovary stained with 1B1 (red), Vasa (blue) and GFP (green) displays aberrant expression of GFP from the GSCs to the 4-cell cyst (100% from GSCs to 2-cell cyst, 13% in 4-cell cyst, n= 25 germaria). GFP channel shown in gray scale in C1. (D, D1) The germarium of a *pgcGFP; twin*^*ry3/ry5*^ mutant ovary stained with 1B1 (red), Vasa (blue) and GFP (green) shows aberrant expression of GFP from the GSCs to the 8-cell cyst (100% from GSC to 4-cell cyst, 40% in 8-cell cyst, n= 25 germaria). GFP channel shown in gray scale in D1. (E) A developmental profile of GFP expression in all stages throughout oogenesis in *pgcGFP, pgcGFP; pum^ET1/FC8^*/*pum*RNAi*, pgcGFP; nos^RC/BN^*/*nos*RNAi and *pgcGFP; twin^ry3/ry5^*/*twin*RNAi ovarioles shows that GFP regulation is lost during the earliest stages of oogenesis in the absence of Pum and its co-factors. (F) PAT assay analysis of *pgc* poly(A)-tail length in GSC tumors and in GSC tumors lacking Pum and Nos. The absence of Pum and Nos results in a longer *pgc* poly(A)-tail length. Scale bars: 10μm.

To test if Pum and Nos control translation of *pgc* mRNA by shortening poly(A)-tail length, as would be expected given the CCR4-Not complex’s role in deadenylation, we utilized the poly(A)-tail length (PAT) assay (Sallés and Strickland 1999). We performed this assay on RNA extracted from GSC-enriched tumors and GSC tumors depleted of Nos and Pum to eliminate the stage of oogenesis in which *pgc* is translationally repressed. In the absence of these RBPs, we detected an increase in the length of the poly(A)-tail compared to the control (Fig. 3F). Together, these observations suggest that Pum, Nos and Twin are recruited to *pgc’s* 3’UTR to suppress its translation in the GSCs by a mechanism that involves shortening of its poly(A)-tail.

We next asked if this regulation of *pgc* by Pum, Nos and Twin is biologically meaningful. Loss of *pum* and *nos* results in failure to maintain GSCs, and this defect is thought to be the result of dysregulation of differentiation-promoting mRNAs in the GSCs (Forbes and Lehmann 1998; Wang and Lin 2005; Gilboa and Lehmann 2004b; Joly et al. 2013). We have previously shown that *pgc* promotes timely differentiation in the pre-CBs where it is expressed (Flora et al. 2018). Thus, we hypothesized that in *nos*, *pum* and *twin* mutants, Pgc is upregulated in the GSCs, forcing the cells to prematurely differentiate and resulting in a loss of GSCs. To test this hypothesis, we made double mutants of *pgc* with *nos*, *pum* and *twin* respectively. Lowering *pgc* levels in all three mutants significantly increased the number of GSCs being maintained (Supplemental Fig. S3M S). Together, our results suggest that Pgc is translationally repressed by Pum, Nos and Twin in the GSCs to ensure appropriate GSC self-renewal and maintenance.

### Me31B cooperates with the decapping protein dGe-1 and *pgc* 5’UTR to mediate repression in the GSCs and early differentiating cysts

Our results suggest that Pum, Nos and Twin regulate *pgc* translation via a conserved sequence in the *pgc* 3’UTR. However, we also found a requirement for the *pgc* 5’UTR in the regulation of *pgc* in undifferentiated cells (Fig. 1G). How could the 5’UTR and 3’UTR of *pgc* cooperate to mediate repression? It has been shown that recruitment of the CCR4-NOT complex also facilitates the recruitment of the de-capping complex to the 5’UTR of mRNAs (Meyer et al. 2010; Garneau et al. 2007; Behm-Ansmant et al. 2006), and that these two complexes at the 5’ and 3’UTR can be bridged by an RNA helicase, DDX6, or Maternal Expression at 31B (Me31B) (Rouya et al. 2014; Ozgur et al. 2015; Nakamura et al. 2001; Fenger-Grøn et al. 2005). This allows “masking” of the mRNAs, making them inaccessible to the ribosome. We therefore hypothesized that Pum, Nos and Twin at the *pgc* 3’UTR could recruit de-capping complex members, such as EDC4 or *Drosophila* Ge-1 (dGe-1), to the cap at the 5’UTR to promote translational repression by masking through the bridging action of Me31B (Fan et al. 2011; Eulalio et al. 2007).

To test this model, we first asked if Me31B associates with *pgc* mRNA. We used a Me31B protein-GFP trap construct and carried out an IP experiment with both anti-GFP and anti-IgG antibodies, in lysates from wild-type and Me31B-GFP trap transgenic ovaries; thereafter we analyzed *pgc* mRNA association using qRT-PCR. We found that there was a significant enrichment of *pgc* mRNA bound to Me31B-GFP protein when compared to IgG IP from the same lysate sample (Fig. 4A). The levels of enrichment were comparable to those of the positive control, *osk* mRNA, which is known to associate with Me31B. Next, we assayed for *pgc*GFP expression upon germline depletion of *me31B* and *dGe-1* and found a loss of GFP repression from the GSC to the 4-cell cyst stage in the presence of *me31B* RNAi and from the GSC to the 8-cell cyst stage for the *dGe-1* RNAi (Fig. 4B-E, Supplemental Fig. 4A). Our results suggest that *pgc* 5’ and 3’UTRs are bridged by a network of RBPs including Me31B and proteins of the decapping complex such as dGe-1 to prevent its translation.

**Figure 4.**
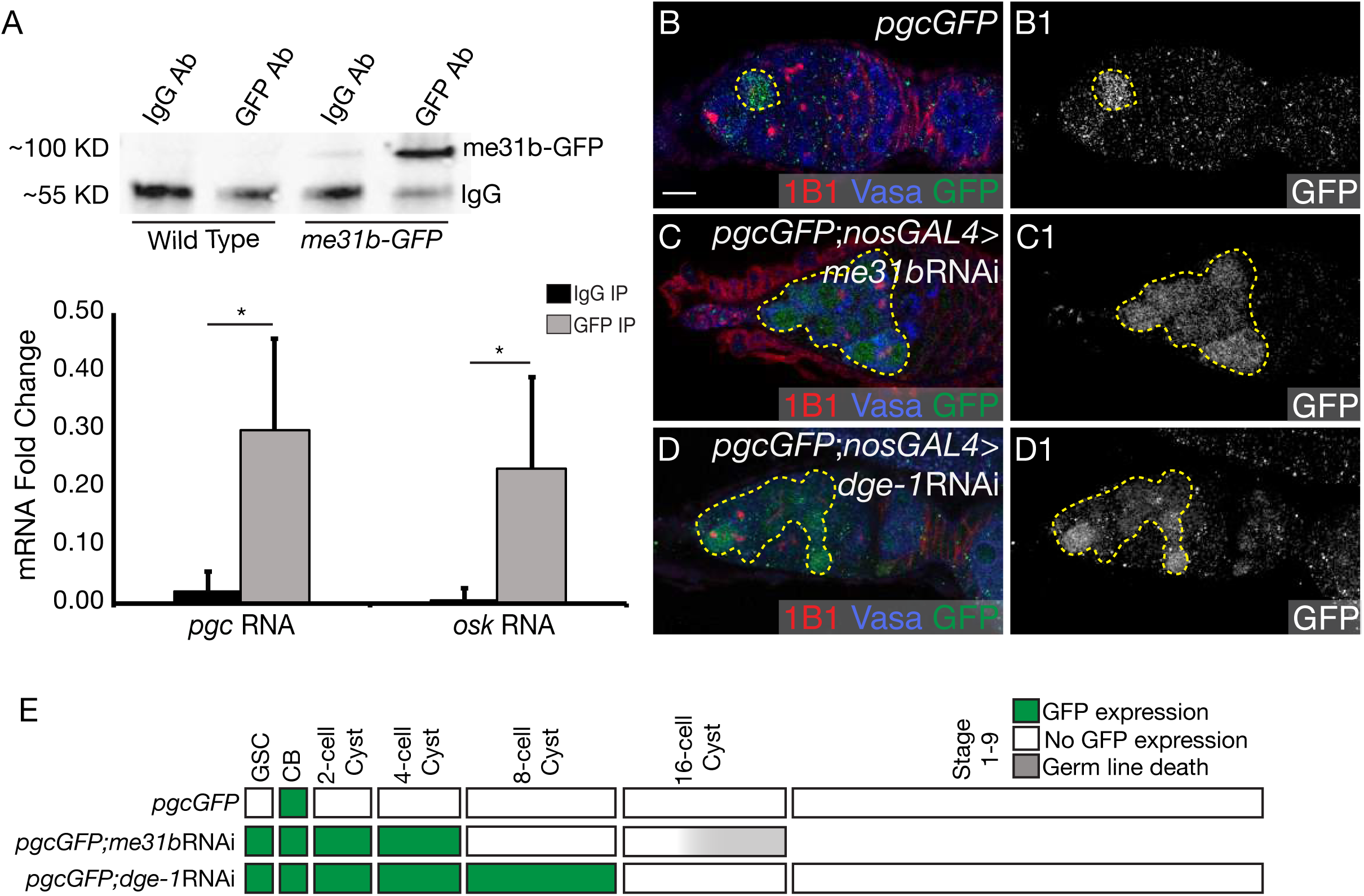
Me31B cooperates with the decapping protein dGe-1 and *pgc* 5’UTR to mediate repression in the GSCs and early differentiating cysts. (A) An IP experiment carried out with GFP antibody in ovary lysates from *me31bGFP*-trap transgenic flies. qRT-PCR analysis of RNA extracted from the IP samples shows that *pgc* RNA is associated with me31b protein *in vivo*, as is the positive control, *osk* RNA. A Student’s t-test statistical analysis was performed. * indicates a p-value <0.05. (B, B1) The germarium of a *pgcGFP* ovary stained with 1B1 (red), Vasa (blue) and GFP (green) shows expression of GFP only in the pre-CB. GFP channel shown in gray scale in A1. (C, C1) The germarium of an ovary with *Me31B* depleted from the germline by RNAi stained with 1B1 (red), Vasa (blue) and GFP (green) shows aberrant expression of GFP in the GSCs to the 4-cell cyst (100% from GSCs to 4-cell cyst, n=20 germaria). GFP channel shown in gray scale in G1. (D, D1) The germarium of an ovary with *ge-1* depleted from the germline by RNAi stained with 1B1 (red), Vasa (blue) and GFP (green) shows aberrant expression of GFP in the GSCs to the 8-cell cyst stages (100% from GSCs to 8-cell cyst, n=20 germaria). GFP channel shown in gray scale in H1. (E) A developmental profile of GFP expression in *pgcGFP, pgcGFP; nosGAL4>me31B*RNAi, and *pgcGFP; nosGAL4>dge-1*RNAi ovarioles shows a temporal loss of GFP regulation restricted to the earliest stages of oogenesis. Scale bar: 10μm.

### Pum and its co-factor Brat regulate Pgc translation in the 4- to 16-cell cysts

Pum can also mediate translational repression via an alternate mechanism by recruiting Brat (Sonoda and Wharton 2001; Muraro et al. 2008; Olesnicky et al. 2012; Harris et al. 2011). Brat engages the cap-binding protein, d4EHP, which competes with the usual cap-binding protein eIF4E, to prevent translational initiation (Cho et al. 2005). Pum is present from the GSCs until the 8-cell cyst and is attenuated from the 16-cell cyst stage onward while Brat is expressed only after the CB differentiates and persists throughout all later cyst stages (Carreira-Rosario et al. 2016; Harris et al. 2011). To test if Pum regulates *pgc* via Brat, we assayed for *pgcGFP* expression in the *pum*^680^ mutant, a separation-of-function mutant that disrupts the interaction between Pum and Brat without affecting the interaction between Pum and Nos (Wharton et al. 1998; Sonoda and Wharton 1999). We found that in *pum*^680^ mutants, there was ectopic *pgcGFP* reporter expression from 4- to 16-cell cyst but not in the earlier stages (Fig. 5A-B1, E, Supplemental Fig. S5A). This observation suggested that Pum may be interacting with Brat and its partner d4EHP to repress *pgc* translation in the differentiating cysts. This to test this, we depleted *brat* and *d4EHP* in the germ line using RNAi. We observed that loss of Brat and d4EHP also results in ectopic expression of GFP from 4- to 16-cell cyst but not in the earlier stages (Fig. 5C-E, Supplemental Fig. S5A). To determine whether this mode of regulation affected the poly(A)-tail length of *pgc*, we performed a PAT assay on *pgc* RNA in *pum*^*680*^ mutants and germline depletions of *brat* and *d4EHP.* We observed no significant change in *pgc* poly(A)-tail length in *pum*^*680*^ mutants and upon depletion of *d4EHP* and *brat* when compared to the control (Supplemental Fig. S5B). A developmental profile of GFP expression in *pgcGFP, pgcGFP; pum^680^, pgcGFP; nosGAL4>brat*RNAi and *pgcGFP; nosGAL4>d4EHP*RNAi shows that compared to the control, loss of Brat and d4EHP results in the loss *pgcGFP* regulation restricted from the 4- to 16-cell cysts (Fig. 5E). These results suggest that Pum not only switches binding partners but also the mode of regulation from a poly(A)-tail dependent mechanism to cap dependent mechanism to regulate *pgc* translation pre- and post-differentiation, respectively.

**Figure 5.**
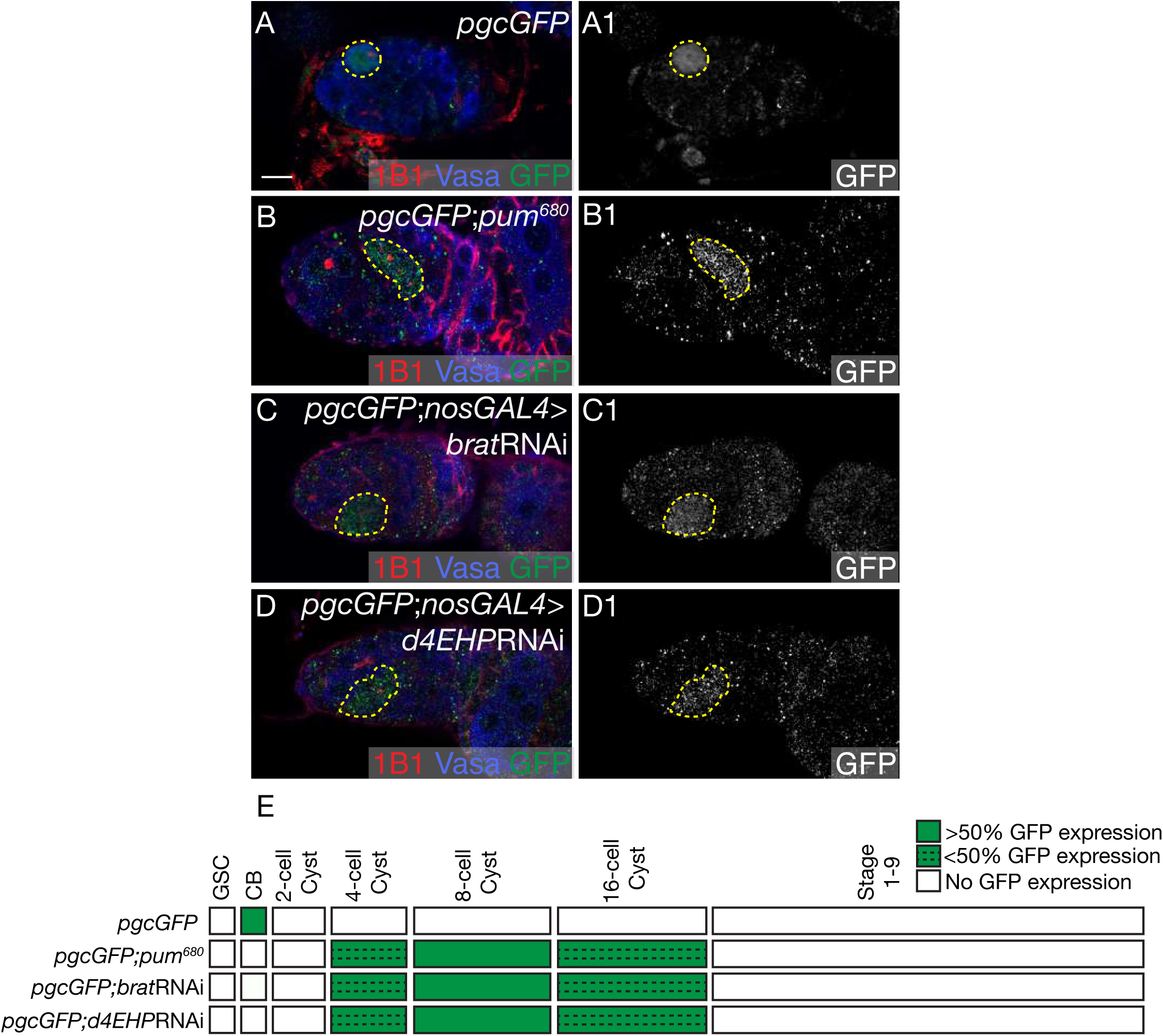
Pum and its co-factor Brat regulate Pgc translation in the 4- to 16-cell cysts. (A, A1) The germarium of a *pgcGFP* ovary stained with 1B1 (red), Vasa (blue) and GFP (green) shows expression of GFP only in the pre-CB. GFP channel shown in gray scale in A1. (B, B1) The germarium of a *pum*^*680*^ mutant ovary stained with 1B1 (red), Vasa (blue) and GFP (green) shows aberrant expression of GFP in the differentiating cysts (25% in the 4-cell cyst, 75% in the 8-cell cyst and 10% in the 16-cell cyst, n=20 germaria). GFP channel showed in gray scale in B1. (C, C1) The germarium of an ovary with *brat* depleted from the germline by RNAi stained with 1B1 (red), Vasa (blue) and GFP (green) shows aberrant expression of GFP in the differentiating cysts (38% in the 4-cell cyst, 54% in the 8-cell cyst and 18% in the 16-cell cysts, n=30 germaria). GFP channel shown in gray scale in C1. (D, D1) The germarium of an ovary with *d4EHP* depleted from the germline by RNAi stained with 1B1 (red), Vasa (blue) and GFP (green) shows aberrant expression of GFP in the differentiating cyst stages (34% in the 4-cell cyst, 62% in the 8- cell cyst and 15% in the 16-cell cyst, n=32 germaria). GFP channel shown in gray scale in D1. (E) A developmental profile of GFP expression in *pgcGFP, pgcGFP; pum^680^, pgcGFP; nosGAL4>brat*RNAi, and *pgcGFP; nosGAL4>d4EHP*RNAi ovarioles shows temporal loss of GFP regulation restricted to the 8- and 16-cell cyst stages in the absence of Brat and d4EHP. Scale bar: 10μm.

### Bru and Cup regulate Pgc translation in the later stages of oogenesis

After differentiation, levels of Pum diminish and levels of Bru increase (Fig. S2G-G2’). We have shown that Bru binds to the 10-nt conserved sequence in the 3’UTR that is required for *pgc* translational control throughout oogenesis (Fig. 2C, F). Therefore, we asked if Bru and its binding partner Cup can repress Pgc translation post-differentiation (Nakamura et al. 2004; Chekulaeva et al. 2006; Kim et al. 2015b). Assaying for the *pgc* reporter in both *bru* mutants and germline depletion of Bru via RNAi we found that translation was de-repressed primarily from the 16-cell cyst stage onwards (Fig. 6A-B1, Supplemental Fig. S6A-B). To determine if Bru recruits Cup to mediate this regulation, we depleted *cup* in the germ line via RNAi, and observed similar ectopic expression of GFP from the 16-cell cyst stage (Fig. 6C). A developmental profile of GFP expression in *pgcGFP; nosGAL4, pgcGFP; nosGAL4>bruno*RNAi and *pgcGFP; nosGAL4>cup*RNAi shows that compared to the control, loss of *bru* and *cup* results in the loss *pgcGFP* regulation primarily from the 16-cell cyst stage onwards (Fig. 6D). To test if Bru and Cup’s mode of regulation affected the poly(A)-tail length of *pgc*, we performed a PAT assay on *pgc* RNA in germline depletion of Bru and Cup. We observed that Bru and Cup depletion results in a dramatic increase of *pgc* poly(A)-tail length (Fig. 6E). As loss of components of the CCR4-Not complex do not show loss of translational control in later stages and poly(A)-tail length increase has been shown to be directly correlated to increased translational efficiency (TE) (Eichhorn et al. 2016; Sachs and Wahle 1993), we favor the model that *pgc* is regulated in the differentiated stages by Bru and its binding partner Cup via a cap dependent mechanism that restricts access to both cap and poly-adenylation machinery.

**Figure 6.**
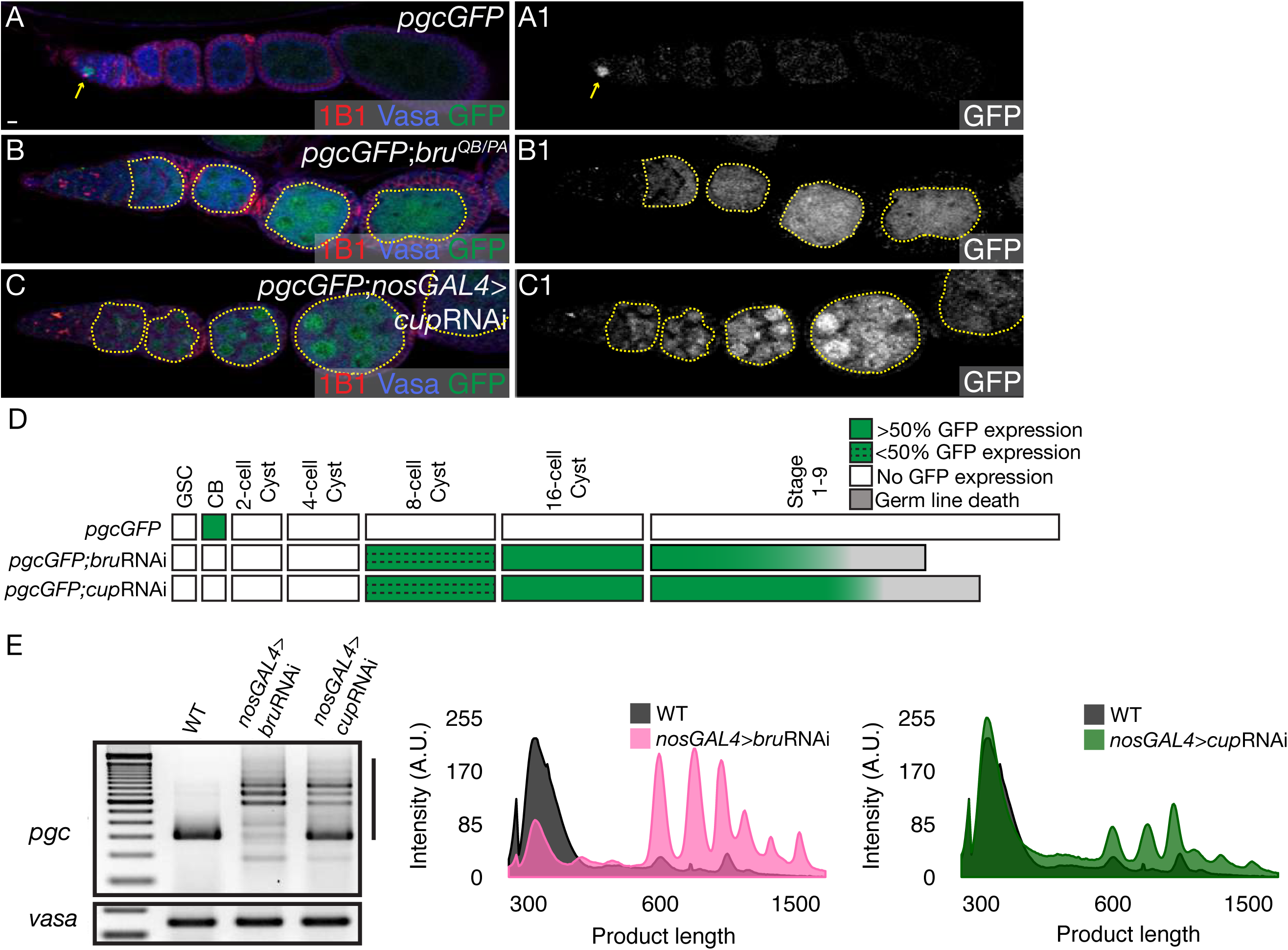
Bru and Cup regulate Pgc translation in the later stages of oogenesis. (A, A1) The ovariole of a *pgcGFP* ovary stained with 1B1 (red), Vasa (blue) and GFP (green) shows expression of GFP only in the pre-CB. GFP channel shown in gray scale in A1. (B, B1) The ovariole of a *pgcGFP; bru*^*QB/PA*^ mutant ovary stained with 1B1 (red), Vasa (blue) and GFP (green) shows aberrant expression of GFP in the entire ovariole beyond the 16- cell cyst stages (12% from 8-cell cyst onwards, 100% from 16-cell cyst onwards, n=25 ovarioles). GFP channel shown in gray scale in B1. (C, C1) The ovariole of an ovary with *cup* depleted from the germline by RNAi stained with 1B1 (red), Vasa (blue) and GFP (green) shows aberrant expression of GFP in the entire ovariole from the later cyst stages (20% from 8-cell cyst onwards, 100% from 16-cell cyst onwards, n=30 ovarioles). GFP channel shown in gray scale in C1. (D) A developmental profile of GFP expression in *pgcGFP, pgcGFP; nosGAL4>bru*RNAi, and *pgcGFP; nosGAL4>cup*RNAi ovarioles shows a temporal loss of GFP regulation throughout oogenesis from beyond the 16-cell cyst stage. (E) PAT assay analysis of *pgc* poly(A)-tail length in ovaries from wild-type, *pgcGFP; nosGAL4>bru*RNAi, and *pgcGFP; nosGAL4>cup*RNAi shows that loss of Bru and Cup in the germ line results in a significant change in the poly(A)-tail length of the *pgc* RNA. Scale bars: 10μm.

### A class of germline RNAs are similarly regulated by both Pum and Bru

Our results show that the conserved RBPs Pum and Bru can recognize and bind the same cis-element in the *pgc* 3’UTR to mediate repression throughout oogenesis. We wondered if this mechanism could be generally applicable for the regulation of maternally deposited mRNAs present in the ovary. To address this, we carried out a Polysome-seq (Poly-seq) experiment that has been successfully used in prior studies to calculate the translational efficiency (TE) of transcripts (Kronja et al. 2014). TE is a measure of actively translating mRNAs, which is achieved by calculating the ratio of mRNA present in the polysome fraction to the mRNA present in the input. Therefore, we utilized this method to identify transcripts that are being inefficiently repressed or being actively translated in the ovaries of *nosGAL4>pum*RNAi and *nosGAL4>bru*RNAi flies when compared to young *nosGAL4* flies. We used young *nosGAL4* ovaries as a control because they do not have mature later stages (stage 10 and onwards) and thus present a similar profile of ovariole stages to those found upon the germline depletion of both Pum and Bru. We conducted RNA-seq of transcripts extracted from the polysome fractions as well as RNA-seq from input RNA and calculated the average TE of all the transcripts in the control and upon germline depletion of Pum and Bru (Supplemental Fig. S7A). We found that when Pum and Bru are depleted in the germline, 1081 and 908 transcripts have higher TE respectively than in the control (Fig. 7A-C). 436 of these transcripts display an increase in TE when either *pum* or *bru* is depleted suggesting that these targets may be co-regulated by them (Fig. 7C). 368 of the 436 transcripts and 179 of the 212 transcripts are maternally provided mRNAs that are also present in mature eggs (Kronja et al. 2014). 212 of the 436 shared transcripts contained a sequence that was similar to the 10-nt PRE/BRE sequence identified in the *pgc* 3’UTR (Supplemental Fig. S7B). Gene Ontology analyses of these 212 shared targets show that these genes are required for gastrulation and cell motility; processes that are mediated by maternally deposited RNAs and occur prior to the maternal-to-zygotic transition of *Drosophila* embryogenesis (Fig. 7D). One such gene that was identified to be co-regulated by Pum and Bru throughout oogenesis was *zelda*, a maternally provided mRNA that plays the role of a master regulator during early *Drosophila* embryogenesis (Fig. 7A-B) (Harrison et al. 2011; Nien et al. 2011; Liang et al. 2008). It is a transcription factor that is required to activate early-developmental somatic genes essential for cellularization, sex determination and body patterning. We do not know if these maternal mRNAs are expressed in the CBs, like *pgc*, or if additional translational regulatory mechanisms silence these mRNAs there. Taken together, our results demonstrate that key determinants for somatic and germ line fate, such as *zelda* and *pgc* respectively are translationally suppressed by Pum and Bru to ensure their repression during oogenesis.

**Figure 7.**
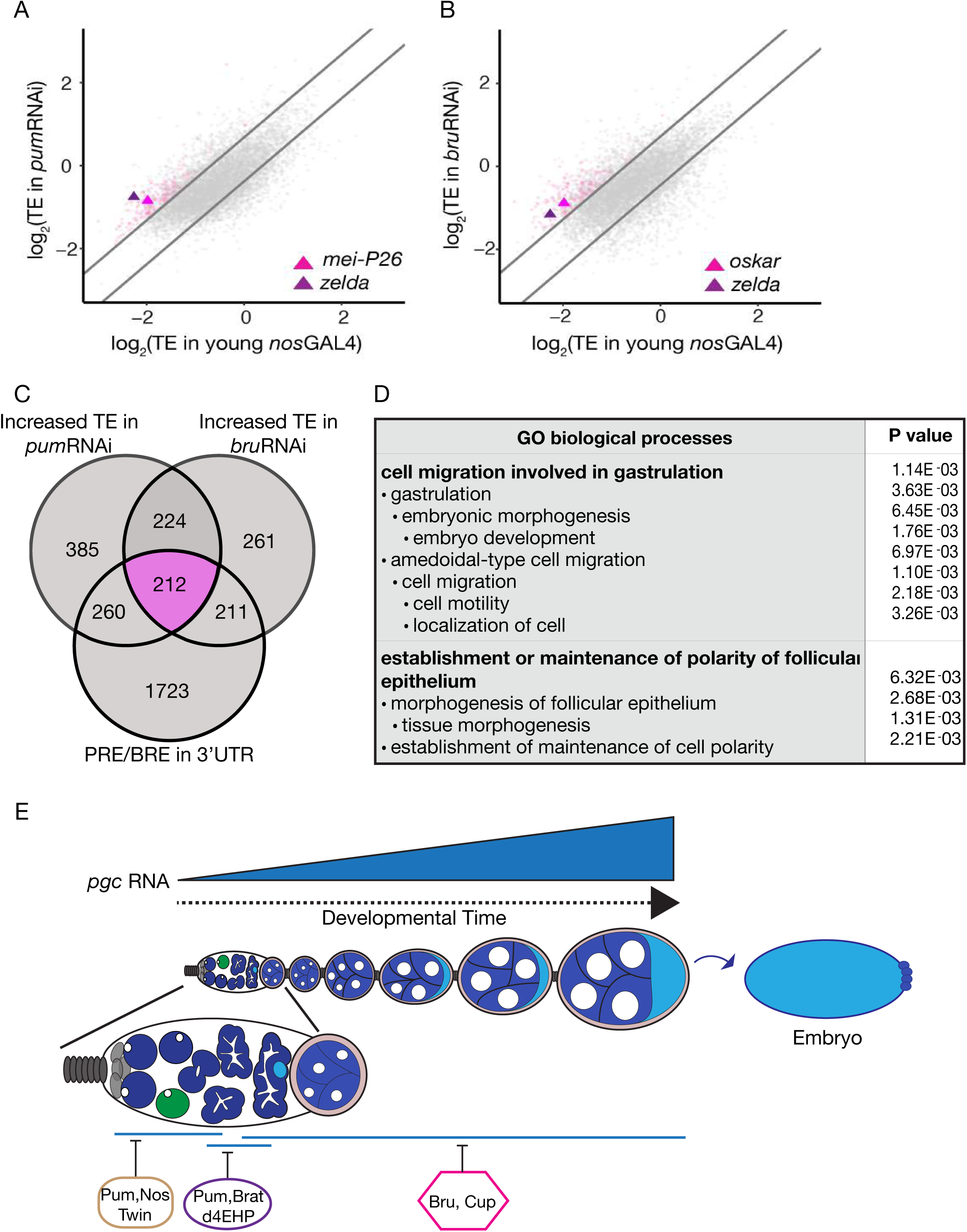
A class of germline RNAs are similarly regulated by both Pum and Bru. (A) A bi-plot representing the translational efficiencies (TEs) of expressed mRNAs in *nosGAL4>pum*RNAi vs young *nosGAL4* (wild-type) ovaries. The lines represent cut-offs, which are one standard deviation above and below the median ratio of TEs. Pink points represent targets containing a PRE/BRE sequence with higher TE in germline depletion of *pum* and *bru*. (B) A bi-plot representing the translational efficiencies (TEs) of expressed mRNAs in *nosGAL4>bru*RNAi vs young wild-type ovaries. The lines represent cut-offs which are one standard deviation above and below the median ratio of TEs. Pink points represent targets containing a PRE/BRE sequence with higher TE upon the germline depletion of *pum* and *bru*. (C) A Venn diagram showing the number of shared targets upon the germline depletion of *pum* and *bru*, which have a higher TE than control and mRNAs that contain an PRE/BRE in their 3’UTR (confusing, reverse the order, first the evidence then the conclusion). The area marked in pink corresponds to the pink points represented in the bi-plots. (D) A table representing the Gene Ontology analysis carried out on the targets of Pum and Bru-mediated regulation that contain a sequence similar to the PRE/BRE sequence identified in the *pgc* 3’UTR. (D) A model accounting for the sequential regulation of *pgc* RNA by various trans-acting factors that are themselves temporally restricted throughout different stages of oogenesis. Pum partners with Nos and Twin to regulate *pgc* in the GSCs to the 8-cell cyst stage. Pum then partners with Brat and d4EHP to regulate *pgc* from the 4- and 16-cell cyst stage. *pgc* is regulated by Bru and Cup from the 8-cell cyst and onwards.

## Discussion

Here we report that a maternal mRNA, *pgc*, is translationally repressed via different temporally restricted RBPs using the same cis-acting sequence during oogenesis. We find that both the *pgc* 5’ and 3’UTRs work in conjunction to regulate translation in the earliest stages of oogenesis. In contrast, during later differentiated stages of oogenesis only the 3’UTR of *pgc* is necessary and sufficient for its translational regulation. We find that a 10-nt conserved sequence in this 3’UTR is essential for *pgc* regulation during the entirety of oogenesis. Surprisingly, two distinct RBPs, Pum and Bru, whose expression is temporally restricted, both recognize and bind this conserved sequence and recruit other cofactors to regulate the mRNA. We find that such regulation is not unique to *pgc*, but that a large class of maternal mRNAs also lose translational control in the absence of both Pum and Bru. Our results indicate that 212 members of this class of mRNAs also share in their 3’UTR a version of the 10-nt conserved sequence necessary for Pum and Bru regulation of *pgc*. These findings suggest that we have identified a broadly utilized mechanism that prevents the translation of specific mRNAs during oogenesis. The fact that some of these mRNAs affect gastrulation and developmental patterning argues that this mechanism evolved to prevent the translation of protein products, which could be deleterious during oogenesis, from mRNAs that must be produced during oogenesis to allow their deposition into the egg to govern the key early steps of embryogenesis.

We find that a dynamic landscape of translational regulators has evolved to allow fine scale control of maternal mRNAs. mRNAs can be regulated either through shortening of the poly(A)-tail mediated by the CCR4-Not complex or through interfering with cap recognition by either the decapping machinery or proteins that bind the cap (Meyer et al. 2010; Garneau et al. 2007; Temme et al. 2014). CCR4-Not complex members as well as decapping machinery proteins are expressed continuously during *Drosophila* germ line development and thus cannot mediate dynamic translational control on their own (Temme et al. 2010; Joly et al. 2013; Fan et al. 2011; Temme et al. 2004). However, carefully choreographed expression of specific RBPs that recognize and bind sequences in the UTRs recruit these regulatory proteins to target transcripts at different stages. Our studies show that Pum, whose expression is restricted to the earliest stages of oogenesis, associates with Nos to recruit the CCR4–Not complex to regulate *pgc* mRNA in the GSCs. After differentiation, Pum switches binding partners and complexes with Brat, a protein only expressed in the differentiating stages, and an adaptor protein, d4EHP, which binds to the mRNA cap to mask *pgc* transcript from the translation initiation factors. As Pum levels diminish, this mode of regulation is handed over to Bru, which is robustly expressed from the 16-cell cyst and onwards, and its partner Cup, which binds to eIF4E to mask *pgc* transcript from the translation initiation factors. Thus, we posit that by utilizing temporally restricted RBPs that can bind the 3’UTR in a combinatorial fashion, the germ line can sculpt differential expression of maternal mRNAs. Surprisingly, we find that for *pgc* mRNA this fine scale translation regulation is mediated by a single conserved sequence in its 3’UTR.

Why does *pgc* use one sequence to bind two trans-acting factors as opposed to utilizing two distinct sequences to bind Pum and Bru independently? Pum recruits Brat, which complexes with d4EHP, that binds the cap to prevent the initiation machinery from accessing the mRNA. Bru recruits Cup which binds eIF4E to prevent the translation initiation machinery from accessing the mRNA. If Pum and Bru are present at the same time, as in the 8-16 cell cyst stage, and bind to different sequences, they will recruit two proteins that have to compete to bind to the mRNA cap. As d4EHP can out compete eIF4E, which is ubiquitously present in all cells, in the presence of Pum, the hand off from Pum to Bru would become difficult. How then is repression of *pgc* mRNA seamlessly handed off from one RBP to another? We observe an overlap for repression mediated by Pum and its two distinct partner complexes in the 4- and 8-cell cysts (Supplemental Fig. S7C). Pum partners with Nos, Twin, Me31B and dGe-1 to repress *pgc* from the GSCs to 8- cell cyst stage while it partners with Brat and d4EHP to regulate *pgc* from the 4- to 16- cell cysts stages (Fig. 7E, Supplemental Fig. S7C). Pum and Bru mediated repression overlap in the 8- and 16-cell cyst stage (Fig. 7E, Supplemental Fig. S7C) We hypothesize that to maintain seamless translational regulation during the 4- to 16- cell cyst stages RBPs compete to bind the same cis-element of their target mRNAs. When levels of one RBP diminish and those of another RBP increase, the RBP present at a lower concentration could be displaced from its binding site on the mRNA, allowing for a smooth transition. Thus, we favor the idea that seamless transitions are mediated by overlapping trans-acting factor regimes and competition for the binding site.

*pgc* is transcribed continuously from GSC stage onwards and accumulates in the oocyte post differentiation. We find that there is a switch in the mode of *pgc* regulation from a Twin (CCR4)-dependent mechanism mediated by Pum which can destabilize mRNAs in the GSCs to a Twin (CCR4)-independent mode mediated by Bru in the later differentiated stages. Loss of Bru during oogenesis results in a dramatic increase in poly-adenylation of the *pgc* mRNA as well as translation of Pgc. This suggests that Bru mediated regulation not only translationally represses *pgc* mRNA during oogenesis but could maintain it in a state poised for poly-adenylation and translation. We also show that this mode of regulation is not unique to *pgc*, and that there is a subset of maternally deposited germ line mRNAs including *zelda* that seem to be regulated similarly. *zelda*, a transcription factor that activates the zygotic genome is expressed at low levels in early embryos and increases as development proceeds concurrent with attenuation of Bru levels (Harrison et al. 2011; Nien et al. 2011; Webster et al. 1997). We hypothesize that post-differentiation it is advantageous to switch the mode of regulation primarily to a cap dependent mechanism mediated by proteins such as Bru to preserve and protect a class of germ line mRNAs that are required to establish the next generation. This guarantees not only seamless translational repression throughout oogenesis, but also serves as an effective strategy to protect and prime these mRNAs to be translated and thus produce the proteins required for early embryonic development.

During mammalian development, maternally synthesized mRNAs are deposited into the egg to support embryonic development and these maternal mRNAs also need to be translationally regulated. Pum and CELF/Bruno-like proteins are both expressed in the mammalian germ line and are required for fertility (Mak et al. 2016; Moore et al. 2003; Mak et al. 2013; Kress et al. 2007). The mammalian homologs of Pum, PUMILIO 1 and 2 (PUM 1 and 2) also bind to a sequence similar to the *Drosophila* NRE, and CELF1/Bruno-like proteins bind to an “EDEN” sequence similar to *Drosophila* BREs (Jenkins et al. 2009; Wang et al. 2001; Vlasova et al. 2008). PUM and CELF/Bruno-like proteins not only play critical roles in the germ line but also required for the development and function of other organs (Siemen et al. 2008; Spassov and Jurecic 2003; Barreau et al. 2006). Both PUM 1, 2 and CELF/Bruno-like proteins are expressed and are required for the proper development of the central nervous system in mice (Meins et al. 2002; Siemen et al. 2011; Wagnon et al. 2011; Zhang et al. 2017). Whether Pum and Bru function together on similar targets in the mammalian germ line and nervous system as they do in the *Drosophila* ovary is not known. Our data suggests that such a hand off mechanism could be acting in these vertebrate systems as well.

---

**Figure S1.**
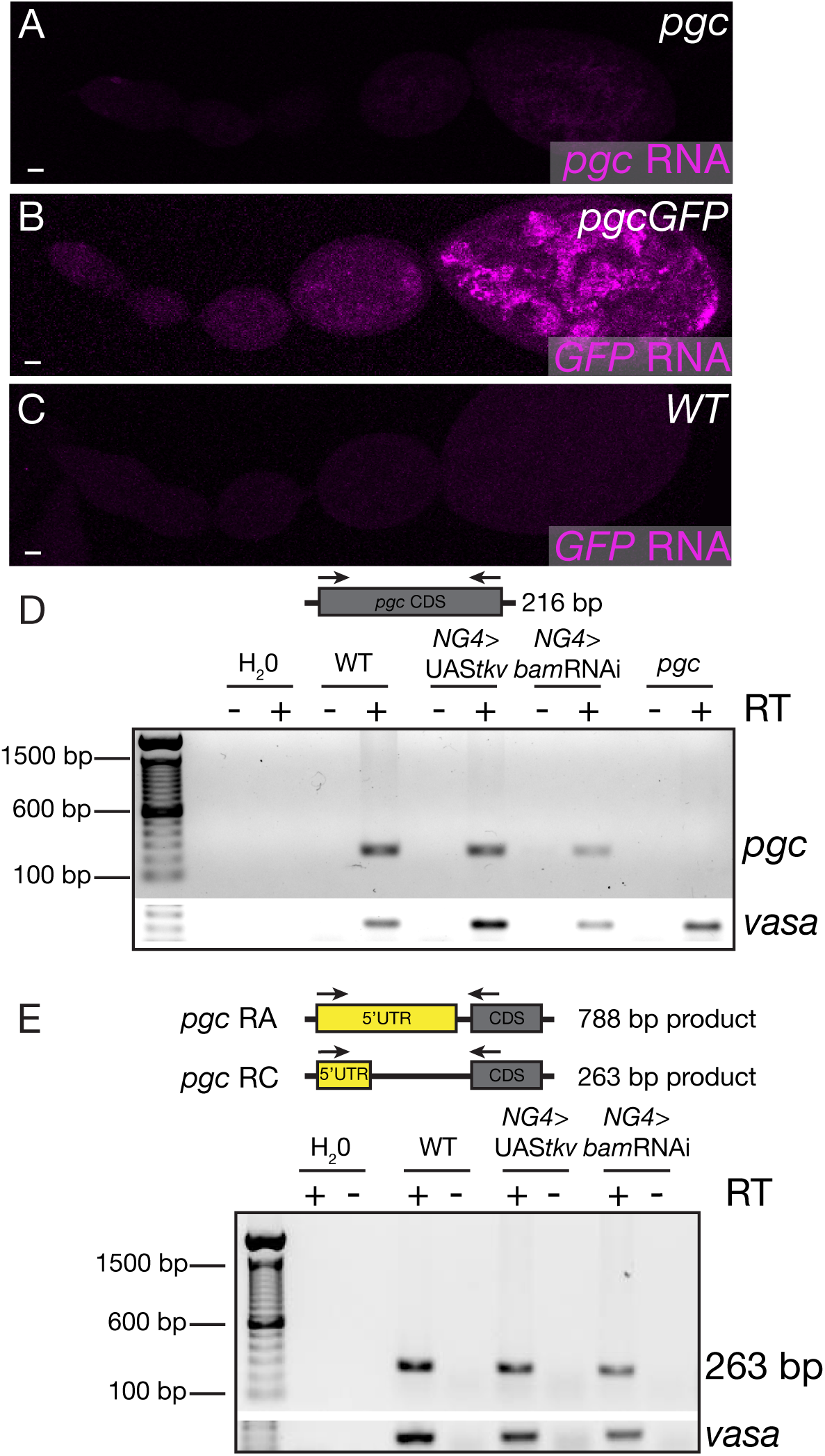
Pgc is translationally regulated via its UTRs. (A) The ovariole of a *pgc* mutant fly probed for *pgc* RNA (magenta) using FISH, show no signal for *pgc* RNA. (B) The ovariole of a *pgcGFP* transgenic fly probed for *GFP* RNA (magenta) using FISH, show similar expression pattern when compared to endogenous *pgc* RNA. (C) The ovariole of a wild-type fly probed for *GFP* RNA (magenta) using FISH, show no signal for *GFP* RNA. (D) RT-PCR of *pgc* CDS was carried out on RNA samples extracted from wild-type, *nosGAL4*>UAS*tkv* and *nosGAL4*>*bam*RNAi show *pgc RNA* is not only present in whole adult ovaries, but also transcribed in GSC and CB enriched tumors. RNA null *pgc* mutant was used as a negative control. RT-PCR of Vasa was carried out as a positive control. (E) RT-PCR of *pgc* 5’UTR was carried out on RNA samples extracted from wild-type, *nosGAL4*>UAS*tkv* and *nosGAL4*>*bam*RNAi. Primers were designed as to show either a 788bp or a 263bp product to confirm what 5’UTR length of *pgc RNA* was being expressed during oogenesis. Results showed presence of short version of *pgc 5’* UTR in whole adult ovaries, GSC and CB enriched tumors. RNA null *pgc* mutant was used as a negative control. RT-PCR of Vasa was carried out as a positive control. Scale bars: 10μm.

**Figure S2.**
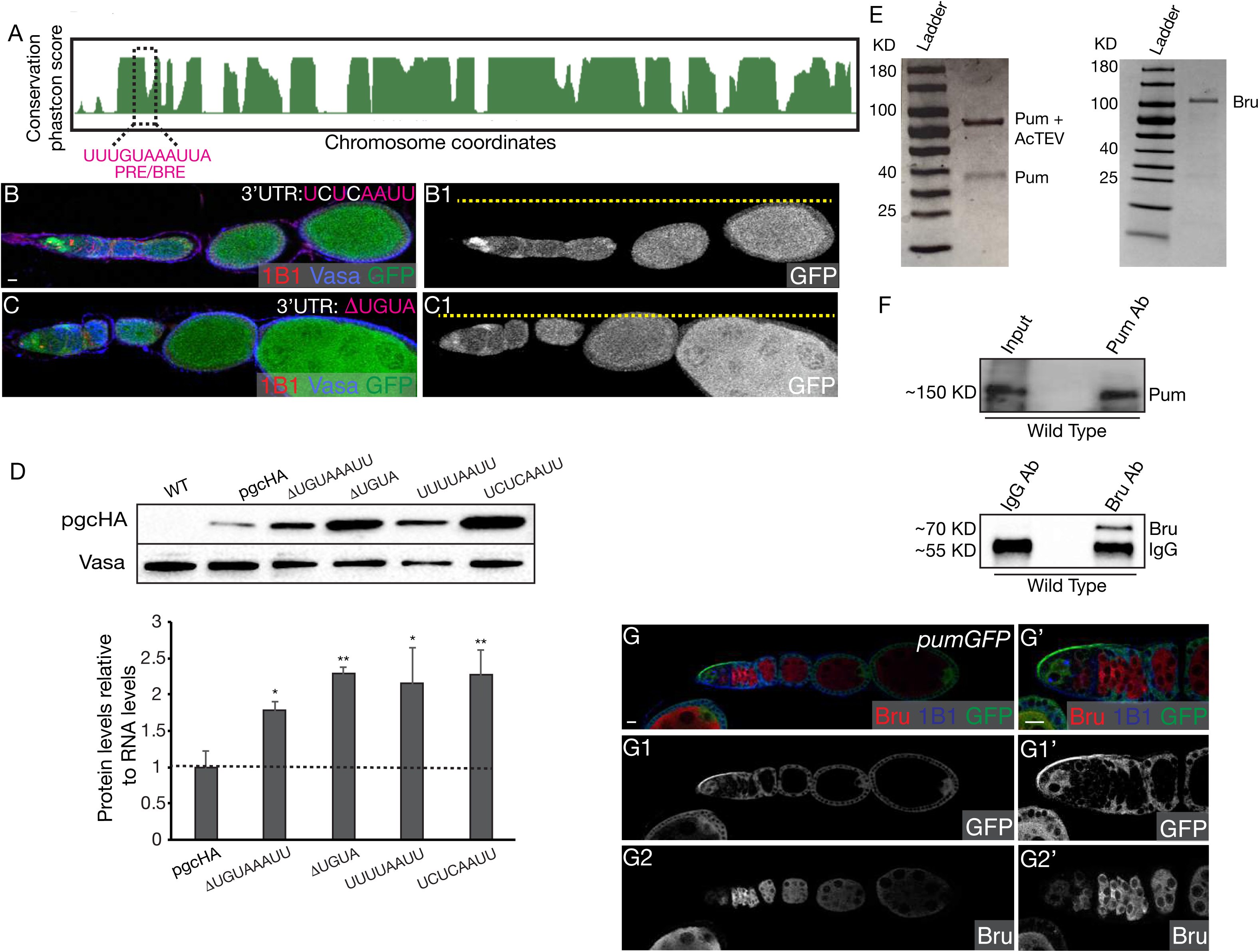
A cis-element in the *pgc* 3’UTR that binds both Pum and Bru is required for translational control throughout oogenesis. (A) A phylogenetic analysis of *pgc* 3’UTR of all Drosophilids identified a conserved sequence that can potentially bind both RBPs, Pum and Bru. (B) The ovariole of a transgenic fly created by fusing GFP to *pgc* 5’ and *pgc* 3’UTR where the UGUA sequence was mutated to UCUC (3’UTR: UCUCAAUU) and driven under *pgc* promoter stained with 1B1 (red), Vasa (blue) and GFP (green) shows loss of GFP regulation throughout oogenesis. (C) Ovariole of a transgenic fly created by fusing GFP to *pgc* 5’ and *pgc* 3’UTR where the UGUA sequence was deleted (3’UTR: ΔUGUA) and driven under *pgc* promoter stained with 1B1 (red), Vasa (blue) and GFP (green) shows loss of GFP regulation throughout oogenesis. (D) Normalized protein expression to RNA levels shows that either deletions or mutations in the PRE/BRE sequence of the 3’UTR of *pgc* results in a significant upregulation of Pgc reporter protein when compared to FL 3’UTR. A student t-test statistical analysis was performed. * indicates p-value <0.05 and ** indicates p-value <0.005. (E) Commasie stained SDS-PAGE gel showing successful purification of recombinant Pum and full length Bru protein. (F) Western Blot shows successful pull-down of Pum and Bru from wild-type ovary lysates using anti-Pum and anti-Bru antibody, respectively. (G-G2’) *pumGFP* transgene fly stained with Bru (red), 1B1 (blue) and GFP (green) shows that Pum protein is expressed in high levels in the earliest stages of oogenesis and lowers in later differentiating stages while Bru protein levels are low in early stages and increases from the 8-cell cyst stages and onwards. G1 and G2 shows GFP and Bru channels in gray. Scale bars: 10μm.

**Figure S3.**
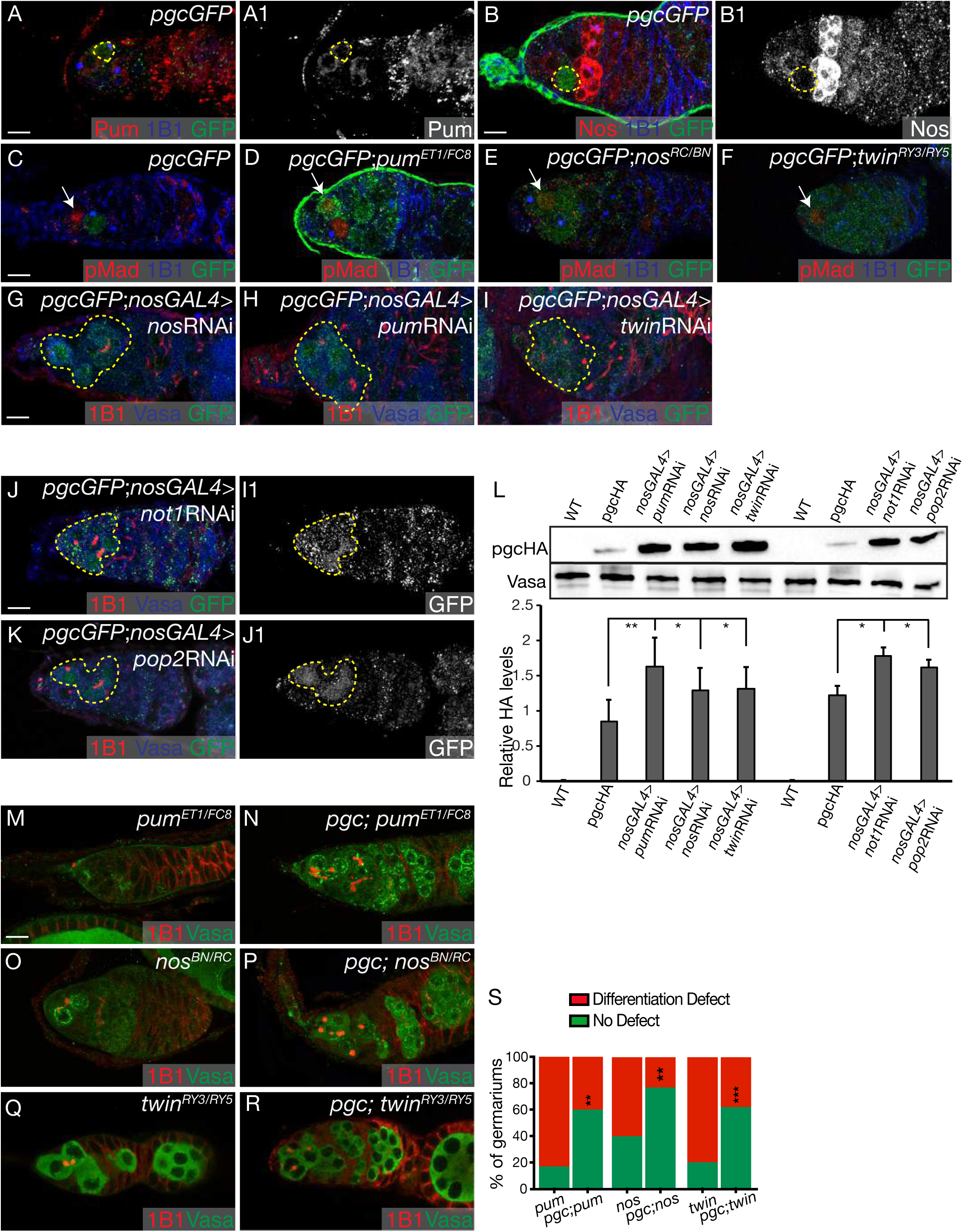
Pum and its co-factor Nos regulate Pgc translation in the GSCs. (A, A1) The germarium of *pgcGFP* fly stained with Pum (red), 1B1 (blue) and GFP (green) shows high levels of Pum protein in the GSC, 2-to 4-cell cysts. Pum staining is shown in gray in A1. (B, B1) The germarium of *pgcGFP* fly stained with Nos (red), 1B1 (blue) and GFP (green) shows Nos protein is present throughout the germarium except for the GFP expressing pre-CB cell. Nos staining is shown in gray in B1. (C) The germarium of *pgcGFP* fly stained with pMad (red), 1B1 (blue) and GFP (green) show GSCs do not express GFP. (D-F) The germaria of *pgcGFP, pgcGFP; pum*^*ET1/FC8*^, *pgcGFP; nos*^*RC/BN*^ and *pgcGFP; twin*^*ry3/ry5*^ stained with pMad (red), 1B1 (blue) and GFP (green) show that in absence of Pum and its co-factors, there is a loss of GFP regulation in the GSCs. (G-I) The germaria of *pgcGFP; nosGAL4>nos*RNAi, *pgcGFP; nosGAL4>pum*RNAi and *pgcGFP; nosGAL4>twin*RNAi flies stained with 1B1 (red), Vasa (blue) and GFP (green) show aberrant expression of GFP in the earliest stages of oogenesis, including the GSCs. (J, J1) The germarium of germline depletion of *not1* ovary stained with 1B1 (red), Vasa (blue) and GFP (green) shows aberrant expression of GFP in the GSCs and 4-cell cysts (100%, n= 25 germaria). GFP channel showed in gray scale in G1. (K, K1) The germarium of germline depletion of *pop2* ovary stained with 1B1 (red), Vasa (blue) and GFP (green) shows aberrant expression of GFP in the GSCs to the 4-cell cyst stages (100%, n= 25 germaria). GFP channel showed in gray scale in H1. (L) A western blot analysis shows a significant upregulation of Pgc reporter protein in the germline depletion of *pum*, *nos*, *twin*, *not1*, and *pop2* ovaries when compared to *pgcGFP.* A student t-test statistical analysis was performed. * indicates p-value <0.05 and ** indicates p-value <0.005. (M, O, Q) The germaria of *pum^ET1/FC8^*, *nos* ^*RC/BN*^ and *twin* ^*RY3/RY5*^ mutants stained with 1B1 (red) and Vasa (green) show germline defects that include proper development of differentiating cysts. (N, P, R) The germaria of *pgc; pum^ET1/FC8^*, *pgc; nos*^*RC/BN*^ and *pgc; twin*^*ry3/ry5*^ double mutants stained with 1B1 (red) and Vasa (green) show rescue of the germline, with proper development of differentiating cysts that eventually make egg chambers. (S) Quantification of rescue experiment shows a significant decrease of differentiation defects in double mutants of *pgc; pum ^ET1/FC8^*, *pgc; nos*^*RC/BN*^ and *pgc; twin*^*ry3/ry5*^ when compared to *pum*, *nos* and *twin* mutants. Scale bars: 10μm.

**Figure S4.**
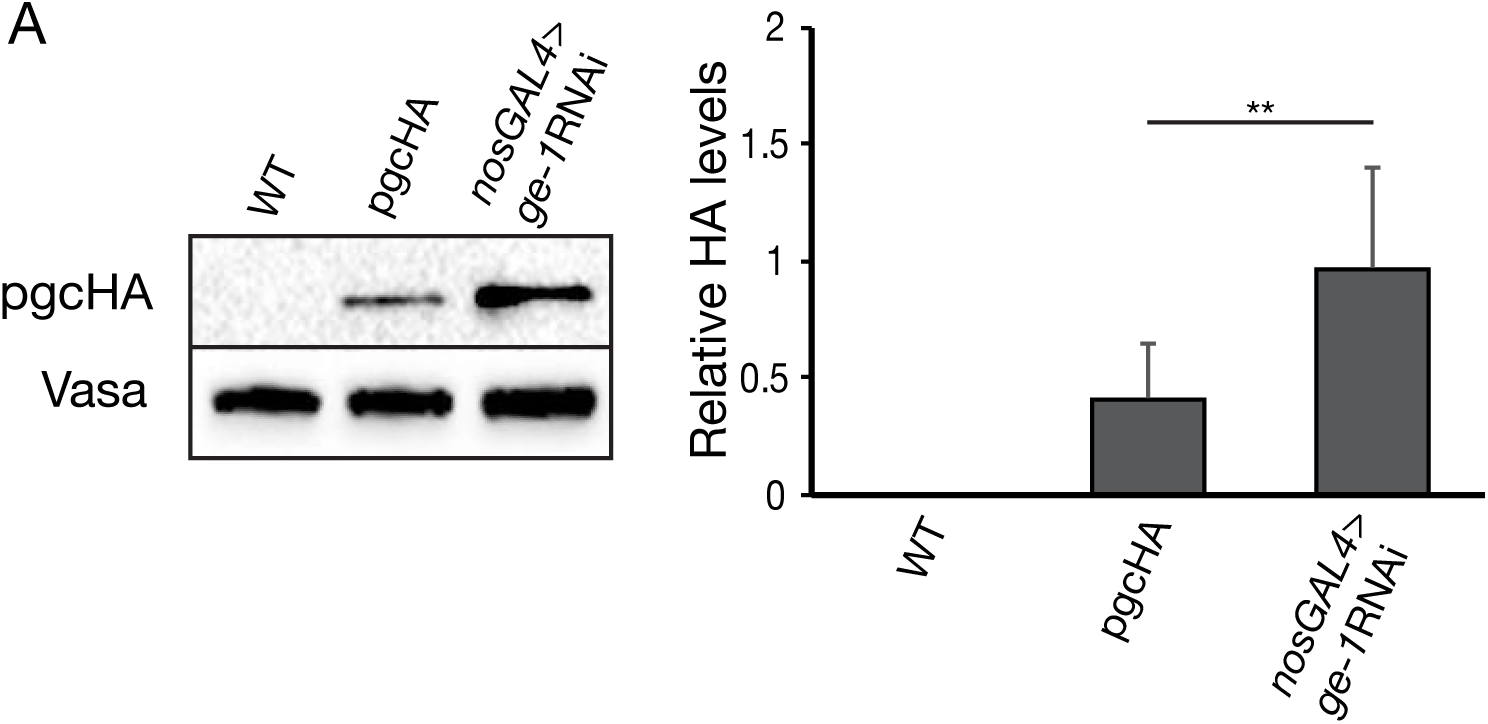
Me31B cooperates with the decapping protein dGe-1 and *pgc* 5’UTR to mediate repression in the GSCs and early differentiating cysts. (A) A western blot analysis shows a significant upregulation of Pgc reporter protein in the germline depletion of *dge-1* ovaries when compared to *pgcGFP.* A student t-test statistical analysis was performed. * indicates p-value <0.05 and ** indicates p-value <0.005. We were unsuccessful in isolating stable lysates from Me31B depleted ovaries to carry out a WB analysis.

**Figure S5.**
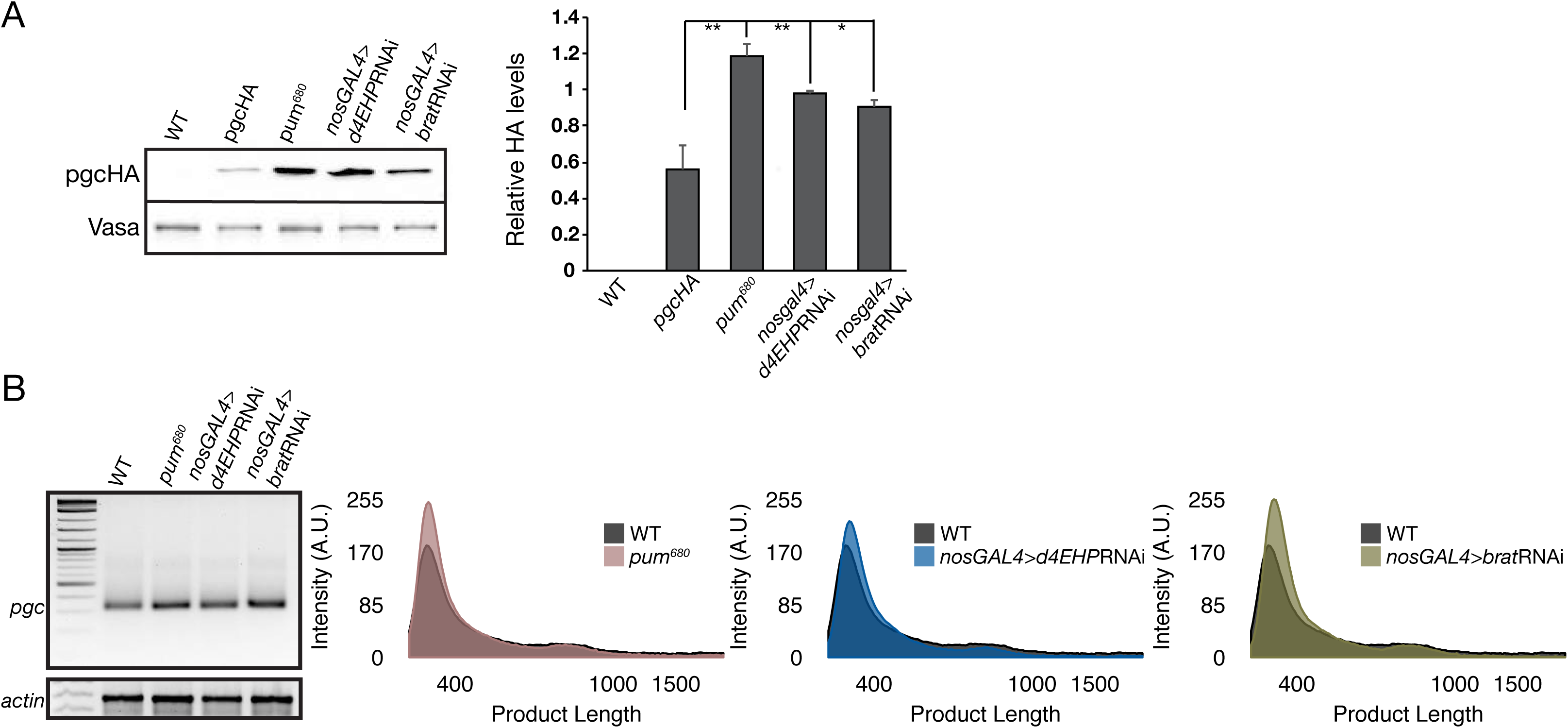
Pum and its co-factor Brat regulate Pgc translation in the 4-to 16-cell cysts. (A) A western blot analysis shows a significant upregulation of Pgc reporter protein in *pum*^*680*^ and the germline depletion of *brat* and *d4EHP* ovaries when compared to *pgcGFP.* A student t-test statistical analysis was performed. ** indicates p-value <0.005. (B) PAT assay analysis of *pgc* poly(A)-tail length in wild-type, *pum*^*680*^ and germline depletions of d4EHP and Brat show that loss of these factors do not result in any change of poly(A)-tail length of *pgc*.

**Figure S6.**
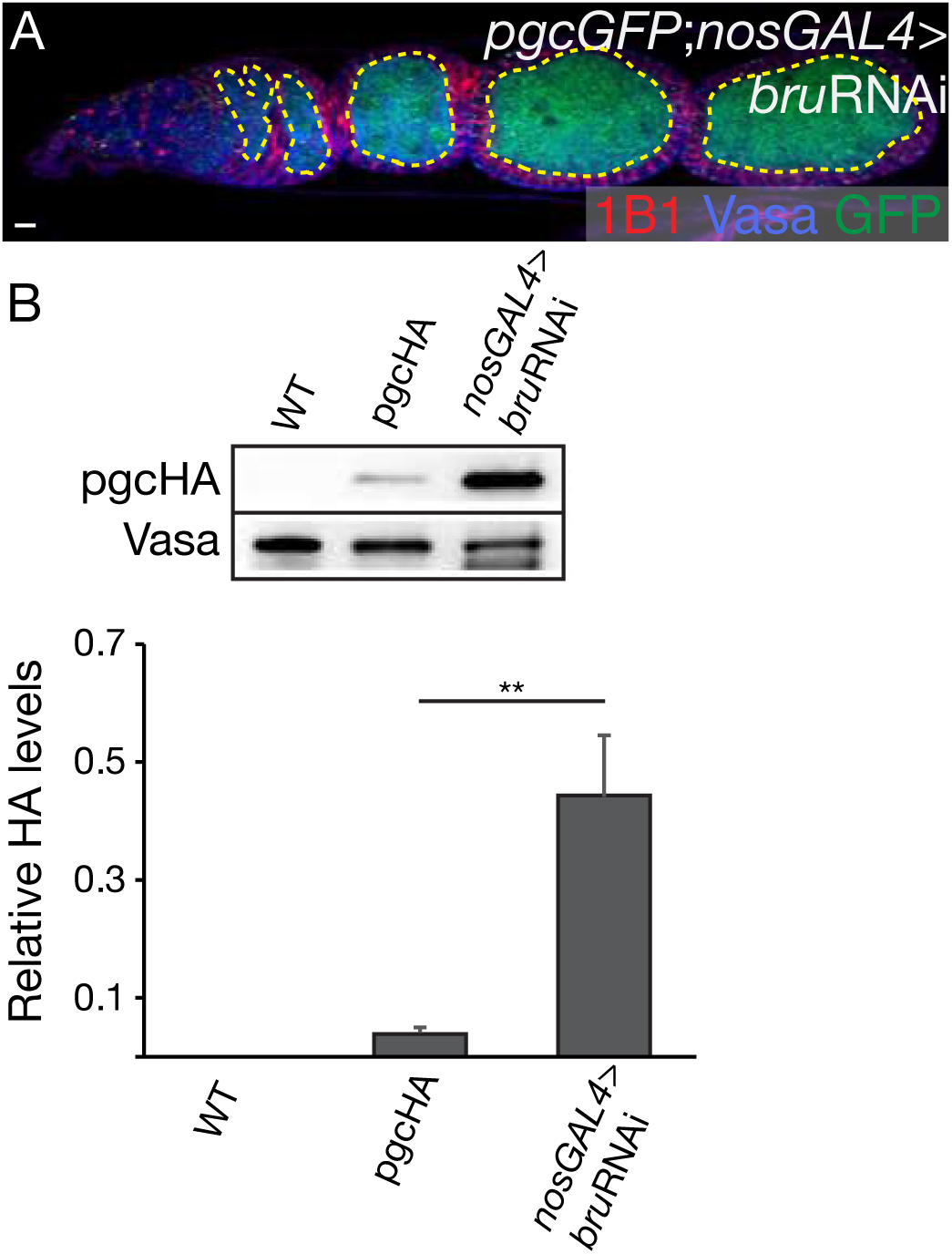
Bru and Cup regulate Pgc translation in the later stages of oogenesis. (A) The ovariole of *pgcGFP; nosGAL4>bru*RNAi stained with 1B1 (red), Vasa (blue) and GFP (green) shows upregulation of reporter expression from 16-cell cyst onwards. (B) A western blot analysis shows a significant upregulation of Pgc reporter protein in the germline depletion of Bru ovaries when compared to *pgcGFP.* We were unsuccessful in isolating stable lysates from Cup depleted ovaries to carry out a WB analysis. A student t-test statistical analysis was performed. ** indicates p-value <0.005. Scale bars: 10μm.

**Figure S7.**
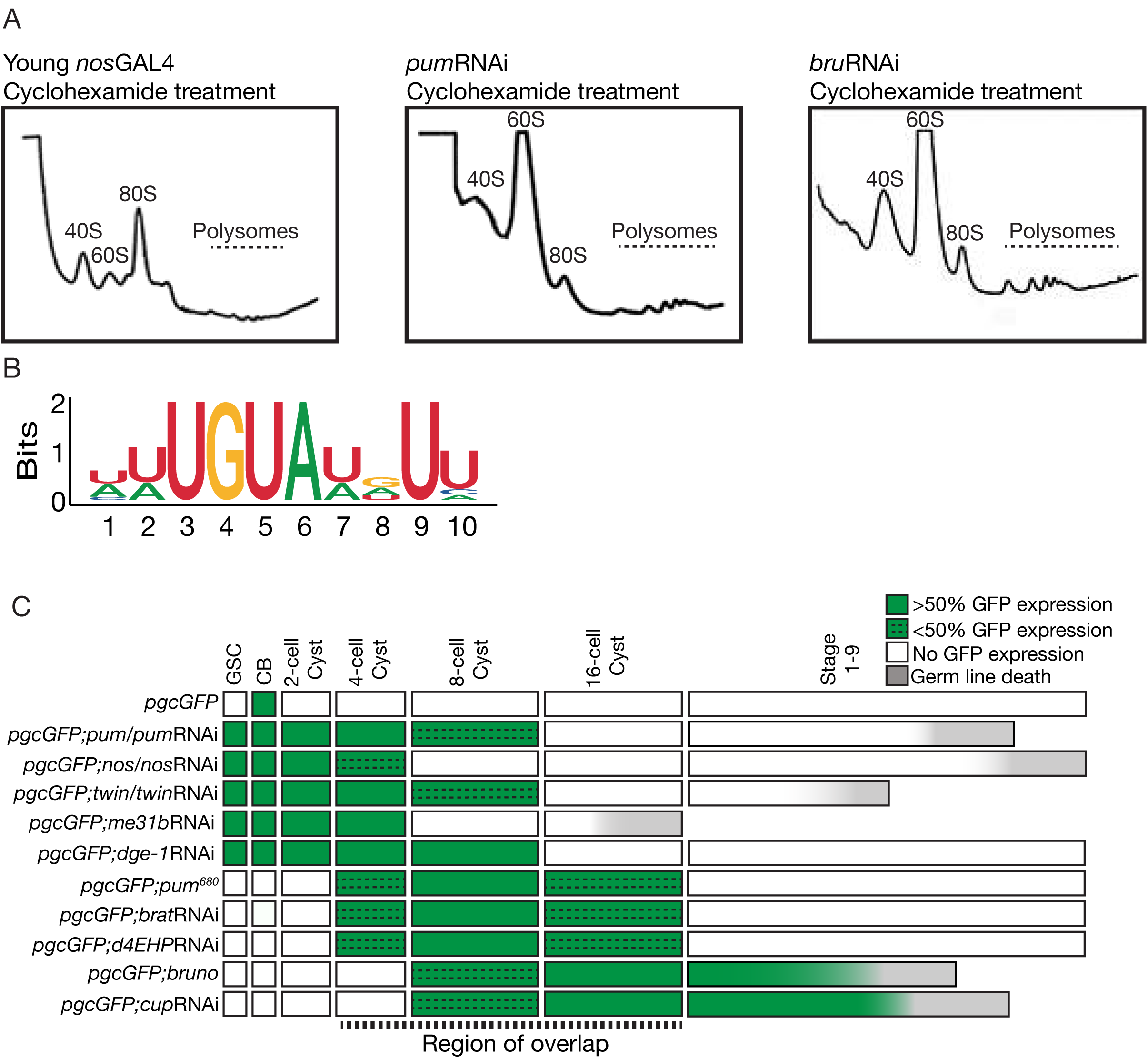
A class of germline RNAs are similarly regulated by both Pum and Bru. (A) Polysome profile traces of young *nosGAL4*, *pgcGFP; nosGAL4>pum*RNAi, and *pgcGFP; nosGAL4>bru*RNAi ovaries treated with cyclohexamide. (B) The logo of the sequences used to identify shared targets of Pum and Bru mediated regulation that contain a sequence similar to the PRE/BRE sequence identified in the *pgc* 3’UTR. (C) A developmental profile of GFP expression in *pgcGFP, pgcGFP; pum^ET1/FC8^*/*pum*RNAi*, pgcGFP; nos^RC/BN^*/*nos*RNAi*, pgcGFP; twin^ry3/ry5^*/*twin*RNAi*, pgcGFP; me31B*RNAi, *pgcGFP; dge-1*RNAi, *pgcGFP; pum^680^, pgcGFP; nosGAL4>brat*RNAi*, pgcGFP; nosGAL4>d4EHP*RNAi*, pgcGFP; nosGAL4>bru*RNAi, and *pgcGFP; nosGAL4>cup*RNAi ovarioles show temporal and sequential loss of GFP regulation in different stages of oogenesis where these trans-acting factors mediate *pgc* regulation.

## Materials and Methods

### Fly stocks

*Drosophila* was grown on corn flour and agar media with brewer’s yeast. All strains were grown at 25**°**C, except RNAi crosses, which were grown at 29**°**C. *pgcGFP and pgc^Δ^* used in this study have been previously reported (Martinho et al. 2004; Flora et al. 2018). *liprinγ^H1^* flies were a gift from the Triesman Lab (Astigarraga et al. 2010). *nos* mutants were generated by crossing the *nos^RC^* and *nos^BN^* alleles (Arrizabalaga and Lehmann 1999). *pum* mutants were created by crossing the *pum*^*FC8*^ and *pum*^*ET1*^ alleles (Forbes and Lehmann 1998). *twin* mutants were created by crossing the t*win*^*ry3*^ and *twin*^*ry5*^ (Morris 2005). The p*um*^*680*^ allele is described in Wharton *et.al*.,1998. *nos-GAL4::VP16* was gifted by the Lehmann lab. *w*^*1118*^, *nos*RNAi (33973, 57700), *pum*RNAi (26725, 38241), *twin*RNAi (32490), *brat*RNAi (28590 and 34646), *d4EHP*RNAi (36876), *not1*RNAi (32836), *pop2*RNAi (30492), *Me31B*RNAi (28566), *ge-1*RNAi (32349), *bru*RNAi (38983) and *cup*RNAi (35406) lines were acquired from the Bloomington Drosophila Stock Center, Bloomington, IN.

### Transgenic lines

The P-P-P (*pgc* promoter-*pgc* 5’UTR-eGFP-*pgc* 3’UTR) construct was generated by cloning eGFP coding sequence into a plasmid with the *pgc* 5’UTR and *pgc* 3’UTR as previously described (Flora et al. 2018).The P-P-T (*pgc* promoter-*pgc* 5’UTR-eGFP-*tubulin* 3’UTR) and P-P-K (*pgc* promoter-*pgc* 5’UTR-eGFP-*K10* 3’UTR) constructs were assembled by PCR amplifying a XhoI-KpnI fragment containing the *α-tubulin* 3’UTR (T) or *K10* 3’UTR (K) was then cloned into the XhoI-KpnI site of the P-P-P plasmid, respectively. In order to allow for interchanging of the 700 bp *pgc* promoter and *pgc* 5’UTR region of P-P-K, AgeI site was created between of those regions of P-P-K via Genscript by Fisher Scientific. The P-N-K (*pgc* promoter-*nos* 5’UTR-eGFP-*K10* 3’UTR) construct was then generated by inserting the *nos* 5’UTR with Agel and Spel overhangs into the AgeI-SpeI site of the P-P-K plasmid. A 700 bp fragment of the *nos* promoter was cloned upstream of the *pgc* 5’UTR of the P-P-K construct at the NotI and AgeI sites to yield N-P-K (*nos* promoter-*pgc* 5’UTR-eGFP-*K10* 3’UTR) construct. The *pgc* 3’UTR fragment was cloned downstream of eGFP at the XhoI-KpnI site of P-N-K to generate P-N-P (*pgc* promoter-*nos* 5’UTR-eGFP-*pgc*3’UTR). The changes to the 3’UTR transgenes in (Fig. 2 and Supplemental Fig. S2) was created by site-directed mutagenesis using Phusion High-Fidelity DNA Polymerase (NEB, Cat # M0530S). The primers used are listed separately.

### Immunostaining

Female Drosophila ovaries were dissected in 1X PBS and fixed in 4% paraformaldehyde for 30 minutes. 1 ml of permeabilization solution, PBST (1X PBS, 0.2% Tween and 1% Triton-X), was added to the tissue. After permeabilization the tissues were blocked in 1 ml of BBT (0.5% BSA in PBST). Then 0.5 ml of primary antibody was added and tissues were incubated at 4**°**C overnight on a nutator. Concentration used for each antibody has been detailed below. After overnight incubation, ovaries were washed three times in 1 ml of BBT for 10, 15, 30 minutes. An additional wash for 30 minutes was carried on by adding 2% Donkey serum to 1 ml of BBT. After the last wash secondary antibody in 0.5 ml of BBT with 4% Donkey serum was added and incubated for 2 hours protected from light. Secondary antibodies used in this study have also been listed below. After the 2-hour incubation, ovaries were washed in 1 ml of PBST for five times. After the washed, one-drop of Vectashield (Vector Labs, Inc.) was added and then the tissue was mounted on a glass slide and a coverslip was placed on the slide. Antibodies used in this study, rabbit anti-Vasa (1:4000 dilution) and chicken anti-Vasa (1:500 dilution) was generated in our lab. mouse anti-1B1 (1:20) is from DSHB, Iowa city, IA. Rabbit anti-GFP (ab6556) (1:2000) and rabbit anti-pSmad3 (ab52903) (1:150) were acquired from abcam, Cambridge MA. Rabbit anti-Nanos (1:500) antibody was a gift from the Buszczak lab. Rabbit anti-Bruno (1:500) and rabbit anti-Pumilio (1:500) antibodies were a gift from the Lehmann lab. Alexa 488 (Molecular Probes), Cy3 and Cy5 (Jackson Labs) conjugated secondary antibodies were used at a concentration of 1:500.

### Fluorescent *in situ* hybridization (FISH)

FISH of the ovaries was carried out probes against *pgc* and *GFP*, which were a gift from the Lehmann lab (Trcek et al. 2017). The ovaries were dissected in 1XPBS, fixed in 3% formaldehyde in PBS for 20 minutes and washed 3 times with PBST. Next, they were treated with 3 ug/ml Proteinase K in PBS and placed on a nutator for 13 minutes at RT, and then placed on ice for 30 minutes. The tissue was then blocked in 2 mg/ml glycine in PBST twice for 10 minutes each and rinsed twice with PBST for 2 minutes. The ovaries were post-fixed for 20 minutes in 3%. The tissue was then washed with PBST 5 times for 2 minutes and washed with pre-warmed fresh pre-hybridization mix (10% deionized formamide in 2X SSC) for 10 minutes. 60 μl per sample of hybridization mix (10% deionized formamide, 0.5 μl of yeast t-RNA, 0.5 μl of salmon sperm DNA, 1 μM of probe, 10% Dextran sulphate, 2 mg/ml BSA, 2X SSC and 1 μl of RNase Out) was added and the sample was incubated overnight at 37°C for at least 12 hours and no more than 16 hours. After incubation, 1 ml of pre-warmed pre-hybridization solution was added to the tissues. After 10 minutes, the pre-hybridization solution was removed, and the ovaries were washed 5 times with 1XPBS for 15 minutes each. After the last wash, PBS was aspirated out and a drop of Vectashield (Vector Labs, Inc.) was added to the tissue before preparing the slide.

### Imaging

All images were taken on a Carl Zeiss 710 Meta confocal microscope using 20X or 40X oil immersion objectives. Scale bars were added using Zen Blue image processing software.

### Western Blot

Twenty wild-type size ovaries or 40 mutant size ovaries were dissected in 1XPBS. After dissection, all the PBS was aspirated and 30 μl of NP-40 buffer with protease inhibitors added to the tissue and homogenized. The lysate was centrifuged at 13,000 rpm for 15 minutes at 4°C. The middle layer was transferred into a new tube. 1 μl of the protein extract was used to carry out a Bradford (Bio-Rad, Cat. #500-0205) assay. 25 μg of protein was denatured with 4X Laemmli Sample Buffer (Bio-Rad, Cat. #161-0747) and β-marcepthanol at 95°C for 5 minutes. The samples were loaded in a Mini-PROTEAN TGX 4-20% gradient SDS-PAGE gels (Bio-Rad, Cat. #456-1094) and run at 110V for 1 hour. The proteins were then transferred to a 0.20 μm nitrocellulose membrane at 100V for 1 hour at 4°C. After transfer, the membrane was blocked in 5% milk in PBST for 2 hours at RT. Primary antibody rat anti-HA (Roche Diagnostics, REF 11867423001) (1:3000) prepared in 5% milk in PBST was added to the membrane and incubated at 4°C O/N. The membrane was then rinsed in 0.5% milk in PBST 4-5 times. anti-rat HRP (1:10,000) was prepared in 5% milk in PBST, and was added to the membrane and incubated at RT for 2 hours. The membrane was then rinsed in PBST 4-5 times. Bio-rad chemiluminescence ECL kit was used to image the membrane. The membrane was then stripped using 25 ml of stripping buffer and re-probed for Rb Vasa (1:6000) as a loading control. anti-rabbit HRP was used at 1:10,000 dilution. For Western Blot analysis *pgcHA* levels were normalized to Vasa levels of each genotype. Then the fold change was calculated for each genotype by subtracting fold change of wild-type control from all experimental samples.

### RNA Extraction

Wild-type ovaries were dissected in 1XPBS. After dissection, all the PBS was aspirated and 100 μl of Trizol reagent was added to the tissue. The tissue was homogenized. 900 μl of Trizol was added, mixed and incubated at RT for 3 minutes. After incubation, 200 μl of Chloroform was added to each sample and mixed vigorously and incubated at RT for 5 minutes. The samples were then centrifuged at 13,000 rpm for 20 minutes at 4°C. The aqueous layer was then transferred to a new centrifuge tube. 2 volumes of 100% ethanol, 10% volume 3 M sodium acetate and 0.5 ul of glycol blue was added to the samples and incubated at −20°C for 1 hour. The samples were then centrifuged at 13,000 rpm for 20 minutes at 4°C. The pellet was then washed with 75% ethanol, air-dried and resuspended in RNase free H_2_O. For efficient re-suspension of the isolated nucleic acid, the sample was incubated at 50°C for 10 minutes. The concentration of the isolated RNA was determined using a Nanodrop. 10μg of nucleic acid was then taken and subjected to a DNase treatment using the TURBO DNA-free Kit by Life Technologies (AM1907).

### Immuno-precipitation (IP)

Each IP experiment was carried out in 100 pairs of wild-type ovaries. Ovaries were dissected in 1XPBS. After dissection, PBS was aspirated and 100 μl of RIPA buffer was added to the tissues and homogenized. Another 200 μl of IP lysis buffer (50 mM Tris pH 8.0, 1% Triton X-100, 0.1% sodium deoxycholate, 0.1% SDS, 140 mM NaCl, 1mM EDTA, 1 mM PMSF, 1 protease inhibitor pill) was added to the lysate and mixed well. The lysate was then centrifuged at 13,000 rpm for 20 minutes at 4°C. The supernatant was transferred to a new tube. 100 μl of homogenate from each sample was transferred to fresh centrifuge tubes. The following antibodies were added to the lysate and incubated at 4°C for 3 hours; 2.5 μl of rabbit anti-GFP (listed above), 1.25 μl of ChromePure Rabbit IgG (Jackson ImmunoResearch Labs), 2 μl of rabbit anti-Bru (gift from Dr. Lilly) or 2 μl of rabbit anti-Pum (gift from Lehmann lab). 100 μl of Dynabeads Protein A (Thermo Fisher Scientific) was rinsed three times with 400 μl of 1:10 dilution of protease inhibitor containing NP-40 buffer. After washing, the beads were re-suspended in 100 μl of NP-40 buffer containing protease inhibitors. 25 μl of these beads were added to each GFP and IgG containing lysate samples and incubated overnight at 4°C. After incubation, the beads were washed four times with 1:10 dilution of protease inhibitor containing NP-40 buffer for 1 minute. An additional two washes for 5 minutes were carried out before re-suspending the beads in 20 μl of NP-40 buffer. 10 μl of beads from each of the samples were used to perform a Western Blot analysis to confirm pull-down. The other 10 μl was used to extract RNA to perform qRT-PCR or RT-PCR experiments to show association of RNA with pulled-down protein.

### Protein Purification

Pumilio expression plasmid pFN18K Pum RNA-binding domain (aa 1091-1426) was gifted to us by the Goldstrohm lab. Pumilio was purified following the protocol described in Weidmann et.al, 2016. Bruno expression plasmid pETM-82 was acquired from EMBL (Chekulaeva et al. 2006). 5 ml of Bruno in pETM-82 in BL21(DE3) was grown overnight at 37°C. This culture was added to 1000 ml of LB-Kanamycin media. Cells were shaken at 220 rpm at 37°C for 2-3 hr or until OD600~0.8The culture was then cooled down to 25°C.0.5 mM IPTG was added to induce the cells and shaken at 220 rpm at 25°C for 3 hours. The cells were then centrifuged at 4000xg for 20 minutes at 4°C in 50 ml aliquots. The pellet was re-suspended in 3 ml of re-suspension buffer (20 mM Na phosphate, 50 mM NaCl, 20 mM imidazole, 10 ul of 500 mg/ml pH 7.4) and sonicated at 20% intensity for 20 seconds for 3 times and pulsed for 20 seconds for 3 times using 1/8 probe, making sure the cell suspension is on ice throughout sonication. The suspension was then centrifuged at 10,000xg for 10 minutes for 4°C. Meanwhile, the column (His GraviTrap, GE Cat#11-0033-99) was equilibrated with 10 mL binding buffer (20 mM Na phosphate, 50 mM NaCl, 20 mM imidazole, 10 ul of 500 mg/ml pH 7.4). The supernatant was added to the column and washed with increments of 1 ml, 4 ml and 5 ml of binding buffer. The protein was then eluted using the following washes; twice with 1 ml of elution buffer (1), twice with 1 ml of elution buffer (2) and three times with 1 ml of elution buffer (3).

> Elution Buffer (1): 20 mM NaPO4, 50 mM NaCl, 150 mM imidazole, pH 7.4
>
> Elution Buffer (2): 20 mM NaPO4, 50 mM NaCl, 300 mM imidazole, pH 7.4
>
> Elution Buffer (3): 20 mM NaPO4, 50 mM NaCl, 500 mM imidazole, pH 7.4

The last two fractions contained purified Bruno protein. 100% glycerol was added to the eluted protein for a final glycerol concentration of 20%. The eluted protein sample was de-salted using the PD-10 column (GE #17-0851-01).

### Electrophoretic mobility shift assays (EMSA)

RNA oligonucleotides were end-labeled using T4 Kinase (NEB) with ATP [γ-^32^P]. Excess ATP was eliminated by using G-25 Sephadix Columns (Roche, Cat # 11273990001). All RNA-binding reaction was performed in 1X Binding Buffer (50mM Tris pH 7.5, 150mM NaCl, 2mM DTT, 0.1mg/μl BSA, 0.001% Tween-20, 0.5μl of dIdC, 1μl RNaseOUT and 0.5μl of yeast t-RNA) (Weidmann et al. 2016). RNA and purified protein was incubated for 20 minutes at RT and then ran on an 8% native polyacrylamide TBE gel at 150V for 4 hours at 4°C. The gel was then dried onto Whatmann filter paper and exposed to a phosphor screen overnight. A Typhoon Trio imager was used to image the EMSAs.

### Real Time-PCR (RT-PCR) and quantitative Real Time-PCR (qRT-PCR)

500ng of DNase treated RNA was reverse transcribed using Super Script III (Life Technologies, Catalog Number: 1808051). For RT-PCR experiments, 1-2 μl of cDNA was amplified using 0.5 μl of 10 μM of each reverse and forward primers, 0.5μl of 10μM (d)NTP and 0.125 μl Taq Polymerase and 2.5 μl 10XTaq Polymerase Buffer. The thermal cycling conditions for PCR was 95°C for 30 seconds, 32 cycles of 95°C for 30 seconds, 3°below the T_m_ of the lowest T_m_ primer for 30 seconds, 68°C for 1 minute, and 1 cycle of 68°C for 4 minutes. After PCR, 2.8 μl of Orange-G dye was added to each sample and 10 μl of PCR product was ran on a 1% agarose gel stained with ethidium bromide to visualize bands.

For qRT-PCR experiments, 0.5 μl of cDNA was amplified using 5μl of SYBR green Master Mix, 0.3 μl of 10μM of each reverse and forward primers. The thermal cycling conditions consisted of 50°C for 2 minutes, 95°C for 10 minutes, 40 cycles at 95°C for 15 seconds, and 60°C for 60 seconds. The experiments were carried out in technical triplicate and three biological replicates for each data point. To calculate fold change in mRNA levels to *RP49* mRNA levels, average of the 2^ΔCt for three biological replicates was calculated. Error bars were plotted using Standard deviation of the ratios. P-value was determined by one-tailed equal variance t-test by comparing ratios of mutants vs. wild-type strains. To calculate relative protein levels to mRNA levels, western blot analysis was carried out, and the fold protein change was divided by fold RNA change from qRT-PCR experiment.

### Poly(A) tail length (PAT) Assay

500ng of DNase treated RNA was reverse transcribed using Super Script III (Life Technologies, Cat.# 1808051) but instead of using oligo (dT), 5 μl of anchored Oligo (dT) primer was used for each sample (Rangan et al. 2009a). 2 μl of cDNA was then amplified using 0.5μl of gene specific forward primer, 0.5 μl of anchored Oligo(d)T, 0.5 μl of 10μM dNTP and 0.125 μl Taq Polymerase and 2.5 μl 10XTaq Polymerase Buffer. The thermal cycling conditions for PCR was 95°C for 30 seconds, 30 cycles of 95°C for 30 seconds, 2° below T_m_ of primer for 30 seconds, 65°C for 1.5 minutes, and 1 cycle of 65°C for 4 minutes. After PCR, 2.8 μl of Orange-G dye was added to each sample and 10 μl of PCR product was ran on a 2.5% agarose gel. The gel was post-stained with ethidium bromide for 20 minutes, and then washed three times with H2O prior to imaging.

### RNA sequencing and sample library preparation

Total RNA was extracted with Trizol, treated with Turbo DNase and poly(A)+ RNA was isolated by double selection with poly-dT beads, using ~6µg total RNA, which is then followed by first-and second-strand synthesis. Sequencing libraries were prepared using NEXTflex Rapid Illumina DNA-Seq Library Prep Kit (Bio Scientific). Samples were single-end sequenced on an NextSeq 500. RNA-seq reads were aligned via HISAT2 (Kim et al. 2015a) set to be splice aware to UCSC dm6 release 6.01. Count tables were generated using featureCounts (Liao et al. 2014).

### Polysome profiling, Polysome-seq and Translation Efficiency (TE) Analysis

~80 ovaries were dissected in PBS supplemented with cycloheximide and frozen immediately with liquid nitrogen. Tissue was homogenized in 200 μl of cold lysis buffer consisting of 1x Polysome buffer supplemented with 1% Triton-X and 1 protease inhibitor pill per 10 ml of buffer. The lysate was centrifuged at 15,000 × g at 4°C for 10 minutes. 20% of lysate was kept aside for “Input RNA” libraries. 750μl of cleared lysate was loaded onto 10-50% sucrose gradients (500 mM KCl; 15 mM Tris-HCl, pH 7.5; 15 mM MgCl2; and 100 μg/ml cycloheximide) in Beckman Coulter 9/16×3.5 PA tubes (Cat. #331372). Gradients were centrifuged at 35,000xg using a SW41 rotor for 3 hours at 4°C. Gradients were fractionated on a Brandel flow cell (Model #621140007) at 0.75 mls/min and 750μl was collected for each fraction with the sensitivity settings at 0.5 Abs. RNA was extracted from the fractions using standard acid phenol: chloroform extraction. The RNA pellet was washed with 80% ethanol and then air-dried. After air-drying the pellet was dissolved in 10 μl of nuclease-free water. Turbo DNase treatment and library preparation was carried out as described above.

To determine translation efficiencies (TE), CPMs (counts per million) values were calculated for all polysome-seq libraries. Any transcript having zero reads in any library was discarded from analysis. The log_2_ ratio between the polysome fraction and total mRNA was calculated and averaged between replicates. This ratio represents translation efficiency. Targets were defined as transcripts falling greater or less than one standard deviation from the median of translation efficiency in both RNAi lines compared to control (Kronja et al. 2014). To discover sequences similar to the pgc BRE in the 3’UTR of targets, all annotated 3’UTRs were downloaded from Flybase for all analyzed targets. A list of BREs and PREs that contain the core sequence UGUA were compiled manually through a literature search. Using the R package Biostrings this list was used to generate and apply a position weight matrix (pwm). This pwm was used to score all 10-mers in all of the previously mentioned 3’UTRs. A minimum score of 90% was chosen as a cutoff by manually ensuring that the core sequence UGUA was present in all targets above the cutoff. Targets identified from polysome-seq were subsetted from the list of RNAs containing a pgc-like BRE in their 3’UTR using a custom R script.

### Oligonucleotides used for EMSA

*pgc* 3’UTR PRE sequence: UUUGUAAAUU

*pgc* 3’UTR ΔPRE sequence: UUAUUGUGAUAUUAUAGUUU

*CycB* 3’UTR NRE sequence: UAGACUAU<u>UUGUAAUUU</u>AUAUC

Scramble sequence: UAAUCAAGAUACAUAUAUGC

*osk* 3’UTR BRE sequence: CUUGA<u>AUGUAUGUU</u>AA<u>UUGUAUGUA</u>UUGAUp890

## Primers

***pgc* CDS_F**: 5’-ATGTGCGACTACCAGATGGAG-3’

***pgc* CDS_R**: 5’-TCAGAATCTCCATCTATCCGCGAT-3’

***pgc* 5’UTR_F**: 5’-CAAGAGAACAAGTTGAGCGTGG-3’

***vasa_*F**: 5’-CGCATTGGACGTACAGGTCG-3’

***vasa_*R**: 5’-TCTTCCTCGACATTGGTGGC-3’

***actin* CDS*_*F**: 5’-GTGTGACGAAGAAGTTGCTGC-3’

***actin* CDS*_*R**: 5’-TCAAAGTCGAGGGCAACATAG-3’

**promoter *pgc*_F**: 5’-GCGGCCGCATAAAAGACTCAAGTTGACCGACATCCCCTTCC-3’

**promoter *pgc*_R**: 5’-GCGCCACCGGTACGGATCTTCGTTTAAGATCTGACC-3’

**5’UTR *α-tub*_F**: 5’-GCGCCACCGGTTCATATTCGTTTTACGTTTGTCAAGCC-3’

**5’UTR *α-tub*_R**: 5’-GCGCGACTAGTATTGAGTTTTTATTGGAAGTGTTTCACACGCG-3’

**5’UTR *nos*_F**: 5’-GGCCGACCGGTTTTAGTTGGCGCGTAGCTT-3’

**5’UTR *nos*_R**: 5’-GGCGCACTAGTGGCGAAAATCCGGGTCGA-3’

**5’UTR *pgc*_F**: 5’-ACCGGTTAGTTTAACATTTTTTTTTCTTCAAGAGAACAAGTTGAGCG-3’

**5’UTR *pgc*_R**: 5-GAGCCAACTAGTTGACTCGAGCTGGACCTCCCA-3

**3’UTR *α-tub*_F**: 5’-CCGCGCTCGAGTGAGCGTCACGCCACTTC-3’

**3’UTR *α-tub*_R**: 5’-CCGCGGGTACCCTTATTTCTGACAACACTGAATCTGGCCG-3’

**3’UTR *K10*_F**: 5’-GCGCCCTCGAGTGAGCAGCCAATGCAACCGAATCCG-3’

**3’UTR *K10*_R**: 5’-GACGGGGTACCGTTGCAAATCTCTCTTTATTCTGCGG-3’

**3’UTR *pgc*_F**: 5’-GCGTCCTCGAGTGACTGGACCTCCC-3’

**3’UTR *pgc*_R**: 5’-GGCCGCCGGTACCCACGATTGCGAATCGAAAATATATTTCTATCTATTTTTTGGG-3’

***pgc* PAT primer 1**: 5’-ACCAGCCTTCAGAGGCGATCGTA-3’

***pgc* PAT primer 2**: 5’-ACCAGCCTTCAGAGGCGATCGTA-3’

**Anchored Oligo(d)T**: 5’-GCGAGCTCGGCGCCCGCGTTTTTTTTTTT-3’

***pgc* qPCR_F**: 5’-CCTCGATGGCATCCTACGAC-3’

***pgc* qPCR_R**: 5’-ATCTCCATCTATCCGCGATGAC-3’

***GFP* qPCR_F**: 5’-GCGACACCCTGGTGAACC-3’

***GFP* qPCR_R**: 5’-GATGTGGCGGATCTTGAAG-3’

***osk* qPCR_F**: 5’-CAACGAAAGGGGCGTGGTGCG-3’

***osk* qPCR_R**: 5’-CGCTGCCGACCGATTTTGTTCCAG-3’

***pgc* 3’UTR ΔPRE mutagenesis**: 5’GACCTCCCAAAAGCCAACTTATTGTGATATATAGTTTTAGCAGTTTTAGCAGTTCGTTTGCCAC-3’

***pgc* 3’UTR UUUUAAUU**: 5’-GGA CCT CCC AAA AGC CAA CTT ATT GTG ATA TT<u>T TTT</u> AAT TAT AGT TTT AGC AGT TCG TTT GCC ACA TG-3’

***pgc* 3’UTR UCUCAAUU**: 5’-GGA CCT CCC AAA AGC CAA CTT ATT GTG ATA TT<u>T CTC</u> AAT TAT AGT TTT AGC AGT TCG TTT GCC ACA TG – 3’

***pgc* 3’UTR ΔUGUA**: 5’-GGA CCT CCC AAA AGC CAA CTT ATT GTG ATA TTA ATT ATA GTT TTA GCA GTT CGT TTG CCA CAT G-3’

## Acknowledgements

We would like to thank the following people for gifting us with re-agents: Dr. Buszczak (UT Southwestern), Dr. Lilly (NIH) and Dr. Lehmann (NYUMC) for providing us with Nanos, Bruno (IP experiment) and Bruno (IF experiment) antibodies respectively. Dr. Salz (Case Western) and Dr. Nakamura (RIKEN) for providing us with *pumGFP* and *me31BGFP*-Trap reporter transgenes respectively. Dr. Goldstrohm (University of Minnesota) for gifting us the Pumilio protein expression vector. Dr. Lehmann (NYUMC) for giving us the *pum*^*680*^ mutant fly. Dr. Trcek (NYUMC) and Dr. Lehmann for providing us with the GFP *in situ* probes. This work was initiated when P.R. was in Ruth Lehmann’s lab at NYUMC.

## Author Contributions

P.F., S.W.D., E.T.M., R.J.P. and P.R. designed experiments, analyzed and interpreted data. P.F., S.W.D., R.J.P., M.N., A.O., P.B., D.P. performed experiments. P.F. and P.R. wrote the manuscript, which all authors edited and approved.

